## Supplementary Materials for "Short-range interactions between fibrocytes and CD8^+^ T cells in COPD bronchial inflammatory response"

**This PDF file includes:**

Supplementary Text
Figures. S1 to S14
Tables S1 to S11
Movies S1 to S5

**Other Supplementary Materials for this manuscript include the following:**

Movies S1 to S5

**Supplementary Text**

The mathematical model is fully described below.

Model description

### 1. Model definition

#### 1.1. *Surface of interest*

Fibrocytes and $CD8^+$ T cells are noted respectively F and C. They evolve on a lattice in the x-y plane of dimension 103 × 103 where the area of each square is determined by the size of a cell (Fig. A) and has a side length of 7 µm, which has been calculated from our *in situ* analyses to match the mean cellular area (around 50 µm$^2$). Indeed, a C cell can be approximated by a disk with a diameter of about 8 µm ((Mrass et al., 2017) and our unpublished observations), giving a surface area of 50 µm$^2$. The size is roughly equivalent for a F cell in lung tissue. Thus, the cells are modeled by squares with a side length $x_0$=7 µm, which correspond to the units of the lattice. Each element of the lattice is defined by 2 coordinates, where the point on the upper left (resp. lower right) corner has the coordinates (1; 1) (resp. (103; 103)). The coordinates of the center of the lattice are (52; 52). The geometry of bronchi corresponds to the transverse section of a cylinder, then we model our surface of interest, the lamina propria, by a zone with a crown shape. The internal radius has been calculated from the mean area of the lumen area combined with the epithelium surface (216 567 µm$^2$), i.e. 38 lattice sites; the external radius has been determined from the mean area of the peribronchial surface (396 436 µm$^2$), i.e. 50 lattice sites. Then the lamina propria $L$ is the set of points with coordinates $(i, j)$ such that :

$$38 \leq \hat{d}\ (i,j),(52,52) \leq 50$$

where $\hat{d}$ is the pseudo-distance : $\hat{d}\left((i,j),(i',j')\right) = \left\lfloor\sqrt{(i-i')^2+(j-j')^2}\right\rfloor$

and $\lfloor x \rfloor$ stands for the integer part of the real number $x$. We thus obtain a working surface containing 3 652 lattice sites (potential cells) corresponding to an area of approximately 179 000 $\mu m^2$, which is in agreement with our in *situ* measurements. In other words, the number $|L|$ of elements of $L$ equals 3 652. Reflecting (zero-flux) boundary conditions are imposed at the external and internal borders. On each site, there is at most one cell.

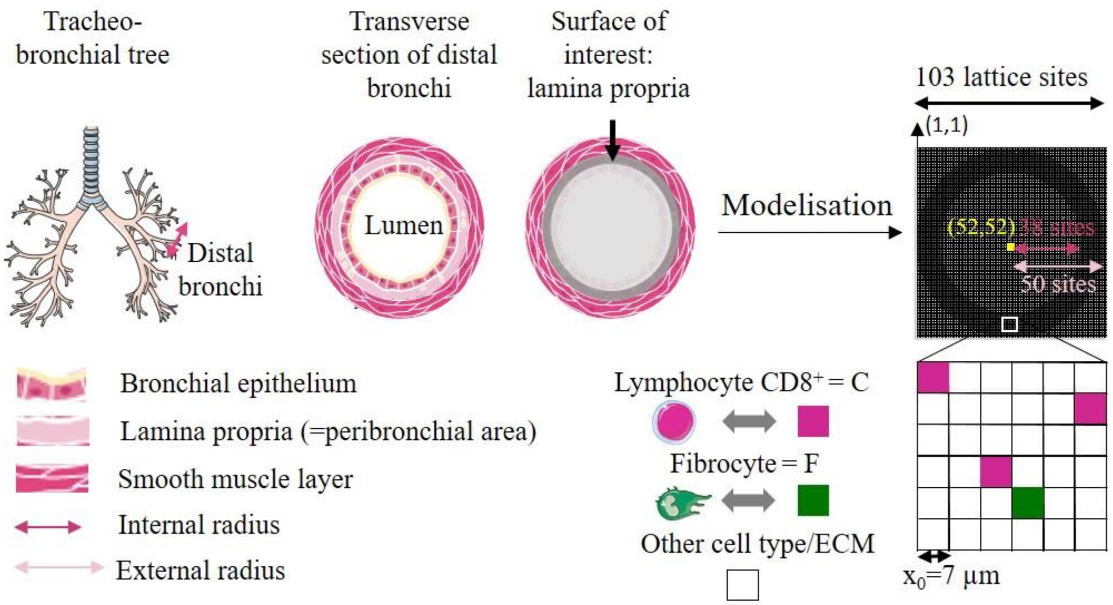


Fig. A. The lamina propria $L$ forms two 2-dimensional crown shape in the bronchial wall, between the bronchial epithelium and the smooth muscle layer. Adapted from (Dupin et al., 2023).

In the literature, it is described that bronchial wall thickness is increased in COPD patients, (Hasegawa et al., 2006; Hogg et al., 2004) but we did not observe this increase in our tissue measurements. We will consider the area of the lamina propria to be the same for healthy subjects and patients with COPD.

1.2. *Neighbourhood*

For any site $(i,j) \in L$, $M(i,j)$ is the neighbourhood of $(i,j)$, it is the set of $(i-1,j-1),(i-1,j),(i-1,j+1),(i,j-1),(i,j+1),(i+1,j-1),(i+1,j),(i+1,j+1)$ belonging to $L$

(Fig. B). For a site inside the lamina propria the cardinal of $M(i,j)$ is 8 and lower if this site is at the edge of $L$. We will note in the following $|M(i,j)|$ the number of elements of $M(i,j)$. In the literature, Moore's neighbourhood is $M(i,j) \cup \{(i,j)\}$.

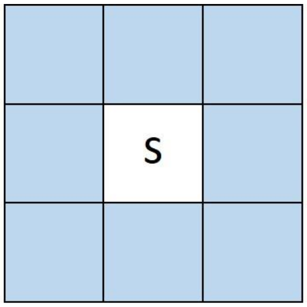


Neighbourhood $M$(s)

Fig. B. Schematic representation showing the neighbourhood $M(s)$ at the site s (shaded blue).

A site of L has the code 1 (resp. 2) if it contains a F (resp. C) cell . If the site is empty it will be coded 0. This state corresponds either to another cell type (mainly mesenchymal, that was hypothetize to interact minimally with fibrocyte and $CD8^+$ T cells) or to extracellular matrix, which again does not play any role in the cellular cross-talk. This is the predominant state, as the bronchial wall contains structural and immune cells sparsely embedded in the extracellular matrix.

A configuration is an element $x = (x(i,j)_{(i,j)\in L}$ where $x(i,j)$ belongs to $\{0,1,2\}$ and $x(i,j) = 1$ (resp. $x(i,j) = 2$) means that a F cell (resp. C) occupies the site $(i,j)$ and $x(i,j) = 0$ when the site $(i,j)$ is empty. The set of configurations is $\{0,1,2\}^L$ and is identified with $\{0,1,2\}^{|L|}$.

For any $s = (i,j)$, $V(F)(s)$ (resp. $V(C)(s)$) denotes the number of F cells (resp. C) near s :

$$V(F)(s) = \sum_{s'\in M(s)} 1_{\{x(s')=1\}},\ V(C)(s) = \sum_{s'\in M(s)} 1_{\{x(s')=2\}}$$

$V(s)$ is the number of F and C cells close to $s$ :

$$V(s) = V(F)(s) + V(C)(s) = \sum_{s'\in M(s)} 1_{\{x(s')=1\ or\ 2\}} = \sum_{s'\in M(s)} 1_{\{x(s')\neq 0\}}$$

1.3. *Initial distribution of cells*

The lattice is initially randomly seeded with both F and C cells at densities described below. For a given patient, $n(F)$ and $n(C)$ are respectively the number of F and C cells divided by the area of the lamina propria (quantified by image analysis). Thus, $n(F)$ and $n(C)$ represent numbers of cells per mm$^2$. We define the initial density $n_0(F+C)$, the initial F cells density $n_0(F)$ and the initial C cells density $n_0(C)$. For the initial distribution, we used the mean distribution of non-smokers subjects, reflecting the "healthy" situation :

$n_0(F+C) = 766$ cells/mm$^2$, *i.e.* 0.766.10$^{-3}$ cells/µm$^2$

$n_0(C)$ = 660 cells/mm$^2$, *i.e.* 0.660.10$^{-3}$ cells/µm$^2$

$n_0(F)$ = 106 cells/mm$^2$, *i.e.* 0.106.10$^{-3}$ cells/µm$^2$

Since the area of the lattice is approximately 179000 µm$^2$, To have an initial density of 0.766.10$^{-3}$ cells/µm$^2$, we will therefore need to add an average value of 0.766.10$^{-3}$ × 179 000 = 137 cells in the grid of 3652 cells. Starting from an initial situation with 137 cells in the grid, this corresponds to an average value of $N_0(C) =$118 C cells and $N_0(F) =$19 F cells.

*1.4 The different time scales*

The median speed of a C cell measured in lung tissue during 15 min is $v_0 =$ 2,3 µm/min (Mrass et al., 2017). For a cell modeled by a square with a side length $x_o =$ 7 µm, $v_0$ represents a movement of one square (lattice square) every 3 minutes. Since we have no information on the in *vivo* speed of a F cell, we will assume that its speed is identical to that of a C cell. Thus, we choose a time step of 3 min for each iteration.

COPD results from a progressive phenomenon: the disease is usually diagnosed starting from the age of 40, but results from exposure to cigarette smoke for several years (Løkke et al., 2006). The time scale that interests us is therefore 10 to 30 years. We thus choose a time period $T$ of 20 years for the simulations. This corresponds to

$$20 \text{ (years)} = 365 \text{ (days)} \times 24 \text{(hours)} \times 20 \text{ (time steps of 3 min)} = 3\,504\,000 \text{ iterations}$$

### 2 The probabilistic interaction model

The mathematical model that we have chosen aims to take into account the interaction between F and C cells and its consequences. We assumed that for a healthy subject as for a patient with COPD, the same model is applied with the same parameters but with different values. These parameters are estimated thanks to experiments and data from the literature and the mathematical results obtained with the streamlined model (Dupin et al., 2023). The notations and parameters of the model are described in Table S10.

First, we first describe the general behavior of a cell, then dissociate the specific cases of F and C. What we call "behavior" includes the description of 4 cellular processes: cell death, displacement, proliferation and infiltration. In a second step, we describe the behavior of all the cells.

#### *2.1 Cell death rules*

C and F cells have a limited lifespan which varies from cell to cell. When they are alive, they will be able to move or duplicate as explained in the following sections. In our algorithm (see section 2.5), when a cell dies, it stays in place for a while and then disappears.

##### 2.1.1 F cell death rules

We define for each F cell a probability $p_{dF}$ of dying (Fig. C).

##### 2.1.2 C cell death rules

For each C cell, we define a "basal" probability $p_{dC}$ of dying, and an increased probability $p_{dC+}$ of dying when the C cell has many other C cells in its neighbourhood. This latter probability is justified as a recent study has shown the existence of CD8$^+$ T cell-population-intrinsic mechanisms regulating cellular behavior, with induction of apoptosis to avoid an excessive increase in T cell population (Zenke et al., 2020). So, we introduce the threshold number σ of neighbouring C cells, above which the probability of dying for a C cell is increased from $p_{dC}$ to $p_{dC+}$.

2 cases are distinguished (Fig. C):

- Case 1: if the C cell has few C neighbors ($V(C)(s)$ <σ, where σ is an unknown integer), then the C cell attemps to die with the probability $p_{dC}$
- Case 2: if the C cell has many C neighbors C ($V(C)(s) \geq$ σ), then the cell C attemps to die with the probability $p_{dC+}$

The numerical values of $p_{dC}$, $p_{dC+}$ and $p_{dF}$ will be presented and justifed in section 3.1.

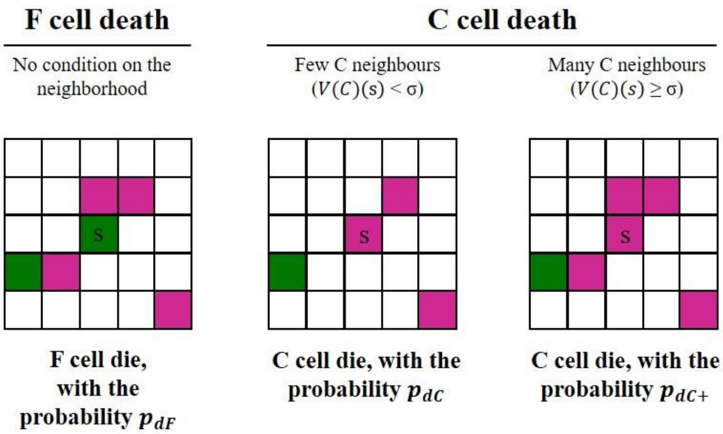


Fig. C. Cell death rules. σ has be taken equal to 3 neighbors. F and C cells are indicated by respectively green and pink squares. Adapted from (Dupin et al., 2023).

#### *2.2 Cell proliferation rules*

Cells have the ability to duplicate, as explained in the following sections. In our algorithm (see section 2.5), when a cell divides, it gives birth to 2 daughter cells, with one staying at the place of the mother cell, and the other one being created in an empty site in the neighbourhood of the mother cell.

#### 2.2.1 F cell proliferation rules

We define for each F cell a probability $p_F$ of dividing.

#### 2.2.2 C cell proliferation rules

For each C cell, we define a "basal" probability $p_C$ of dividing and an increased probability $p_{C/F}$ of dividing when the C cell has F cell(s) in its neighbourhood. This latter probability is justified as our results and those from another study (Afroj et al., 2021) show a robust and high increase of C cell proliferation in direct co-cultures of F and C cells.

Consider a C cell located in $s$. To reflect contact inhibition that enables cells to stop proliferating when many of them are in contact with each other, we also introduced a threshold number λ.

Two cases are distinguished (Fig. D):

- Case 1: if all the sites of $M(s)$ are occupied (Case 1a, *i.e.* $V(s) = |M(s)|$), or if all empty s' sites belonging to $M(s)$ have "many" C neighbours (Case 1b, *i.e.* $V(C)(s') \geq \lambda$, where λ will be taken equal to 3) (Case 1b), the C cell does not divide.
- Case 2: there is at least one empty site s' belonging to $M(s)$ and such as $V(C)(s') < \lambda$. If there is no F cell in $M(s)$, then the C cell attempts to divide with the probability $p_C$ (case 2a). If there is at least one F cell in $M(s)$ ($V(F)(s') \geq 1$), the C cell attempts to divide with the probability $p_{C/F}$ (case 2b).

If the C cell divide, the C cell remains in $s$ and we uniformly choose an unoccupied site $s$' belonging to $M(s)$, such that $V(C)(s') < \lambda$, on which a new C cell is created.

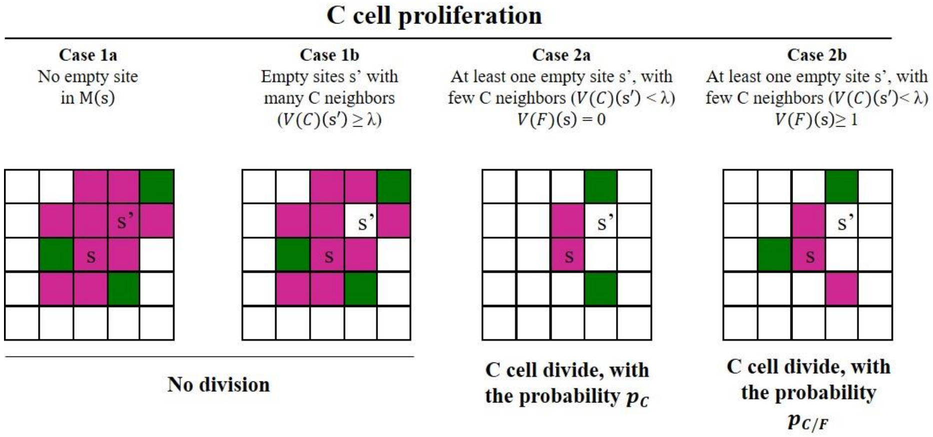


Fig. D. The different cases for C cell proliferation rules. λ has be taken equal to 3 neighbors. F and C cells are indicated by respectively green and pink squares. Adapted from (Dupin et al., 2023).

*2.3 Cell displacement rules*

C and F cells are able to move, as shown previously (Dupin et al., 2016; Mrass et al., 2017). This process is taken into account in the model, as described below.

Let $s = (i, j)$and $s' = (i', j')$ be two sites of the lamina propria. We define for each F cell (or C cell) a probability $P_F\ (s, s')$ (respectively $P_C\ (s, s')$) of going from s to s'. A living cell can only move to a site adjacent to the site it occupies:

$P_F\ (s, s') = P_C\ (s, s') = 0$ if s' $\notin M(s) \cup \{s\}$, or if s' $\in M(s)$ and is occupied.

#### 2.3.1 F cell displacement rules

Our image analysis on lung tissue indicate that (i) the minimum median distance between two F cells is relatively high (30.6 µm for control subjects, 21.5 µm for patients with COPD) and (ii) F cells are rarely observed in clusters of two or more F cells, suggesting that F cells does not attract each other. We will assume that F cell is not attracted by another F cell. Then, we applied $f_F$ to the variable $V(C)(s')$, the number of C cells located in the Moore's neighbourhood of s'.

A living F cell, located in s, will move to the site s' $\in$ M$(s) \cup \{s\}$, with the probability :

$$P_F(s,s') = \begin{cases} k_F\, f_F\left(V(C)(s')\right), & if\ \ s' \in \mathrm{M(s)},\ \ s' \text{ is empty and } s' \neq s \\ k_F\, x_F\,, & if\ s' = s \end{cases}$$

Where :

1. $k_F$ is a constant such that $P_F(s,.)$ is a probability:

$$k_F = \frac{1}{x_F + \sum_{s' \in M(s)} f_F\left(V(C)(s')\right) 1_{\{s' empty\}}}$$

2. $x_F > 0$ will be determined later (see section 3.3.3)
3. $f_F$ is a function which will defined in section 3.3.1

$$f_F(n) = \begin{cases} 1\,, & for\ n\ to\ be\ defined\ (see\ section\ 3.3.1) \\ \varepsilon_F\,, & for\ n\ to\ be\ defined\ (see\ section\ 3.3.1) \end{cases}$$

#### 2.3.2 C cell displacement rules

Our image analysis on lung tissue indicate that (i) the minimum median distance between two C cells is low (7,9 µm for control subjects, 4,9 µm for patients with COPD) and (ii) C cells are often observed in clusters of two or more C cells, suggesting that C cells can attract each others. On the other hand, C cells express CCR1, CCR2, CCR4, CCR5, CXCR1 and CXCR2 (Hombrink et al., 2016), which are receptors of the chemokines CCL2, CCL3, CCL4, CXCL1 and CXCL8 that can be secreted by F cells (Dupin et al., 2018), and the minimum median distance between two C cells is low (10,9 µm for control subjects, 6,8 µm for patients with COPD), indicating that C cells can be attracted by F cells. Therefore, in contrast to $f_F$ which is applied to $V(C)(s')$, we applied $f_C$ to the variable $V(s')$, the total number of C and F cells located in the neighbourhood $M(s')$.

A living C cell, located in $s$, will move to the site $s$' $\in$ M(s) $\cup$ $\{s\}$, with the probability :

$$P_C(s,s') = \begin{cases} k_C\, f_C\left(V(s')\right), & if\ \ s' \in \mathrm{M(s)},\ \ s' \text{ is empty and } s' \neq s \\ k_C\, x_C\,, & if\ s' = s \end{cases}$$

Where :

1. $k_C$ is a constant such that $P_C(s,.)$ is a probability:

$$k_C = \frac{1}{x_C + \sum_{s' \in M(s)} f_C\left(V(s')\right) 1_{\{s' empty\}}}$$

1. $x_C > 0$ will be determined later see section (see section 3.3.3)
2. $f_C$ is a function defined on $\{1, \ldots, 8\}$ will be found in section 3.3.2:

$$f_C(n) = \begin{cases} 1, & for\ n\ to\ be\ defined\ (see\ section\ 3.3.2) \\ \varepsilon_C, & for\ n\ to\ be\ defined\ (see\ section\ 3.3.2) \end{cases}$$

*2.4 Cell infiltration rules*

F and C cells can infiltrate the lungs at stable state, and this process can be amplified during exacerbations. We will add at the beginning of each 3 min period (see section 1.4), one cell F (resp. C) with the probability $p_{istaF}$ (resp. $p_{istaC}$ ) to take into account the phenomenon of infiltration during at the stable state. These probabilities will be determined from biological considerations (see section 3.4). If a cell is recruited, we randomly and uniformly position it among all the empty sites of the lamina propria (Fig. E). If there are no empty sites nearby, no cell is added.

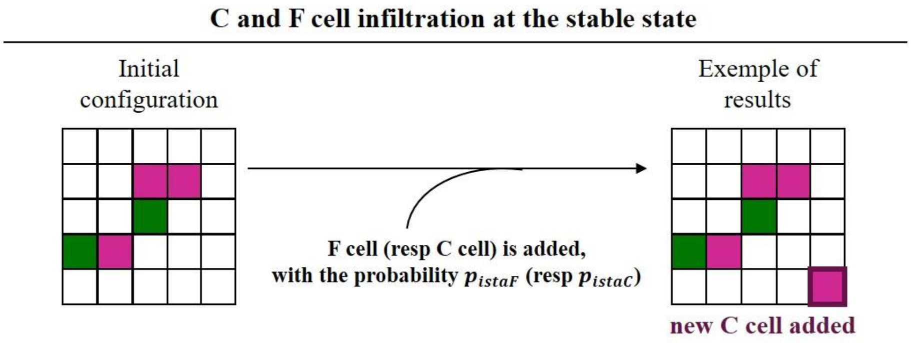


Fig. E. C and F cell infiltration rules at the stable state. F and C cells are indicated by respectively green and purple squares. Adapted from (Dupin et al., 2023).

The process of infiltration can be amplified during exacerbations, which is an acute event specific of patients with COPD, and not happening in healthy subjects. To take into account the infiltration during the exacerbations, we will add every year a number $N_{iexaF}$ (resp. $N_{iexaC}$) of F cells (resp. C) with the probability $p_{iexaC}$ (resp. $p_{iexaF}$). If cells are added, they are placed uniformly on the empty sites of the lamina propria.

Concerning $p_{iexaF}$, it depends closely on the condition of the subject (healthy *vs* COPD). For a healthy subject, as there is no exacerbation, this probability is zero. The values of $p_{iexaF}$ and $N_{iexaF}$ for a patient with COPD will be specified in section 3.4.1.

#### *2.5 Dynamics of C and F cells*

The time steps are denoted $k$, where $1 \leq k \leq T$. The dynamics take place by time step (Fig. F). Let us consider the beginning of the time step $k + 1$. They are $N_k(F)$ F cells and $N_k(C)$ C cells, then $N_k = N_k(F) + N_k(C)$ is the total number of cells in the lamina propria at the beginning of the time step $k$.

If a C or F cell is added by infiltration at the stable state (see section 2.4), it is positioned randomly and uniformly across all vacant sites. This implicitly assumes that all the sites are not occupied. Otherwise, no cell is added. We agree that the added cell is not part of the $N_k$ initial cells, and cannot be drawn at random afterwards. It is therefore neither subject to death, nor to duplication, nor to displacement during the time step $k + 1$, it is just considered as present.

We divide the time step $k + 1$ into $N_k$ sub-time steps. For each sub-time step, we randomly draw one cell among the $N_k$ cells (with the probability $1/N_k$). Several cases can occur.

1. If the selected cell is dead or if it is a C cell that gave birth to a new cell by division in a previous sub-time step, nothing happens.

**2.** Assuming in the following that the selected cell is alive and is not a "mother" cell, we note $(i, j)$ the site occupied by this cell.

a. A F cell attempts to die with the probability $p_{dF}$,

b. A C cell attempts to die with the probability $p_{dC}$ or $p_{dC+}$ according to the rules defined in section 2.1.2.

**3.** We hypothetize in the following the randomly selected cell did not die.

a. A F cell moves according to the procedure described in section 2.3.1.

b. A C cell divides according to the procedure described in section 2.2.2. If the C cell divides, we uniformly choose an empty site *s'* belonging to $\mathrm{M(i, j)}$, such that $V(C)(s') < \lambda$, on which a new C cell is created. The new C cell is not part of the population of $N_k$ cells, it will be added once the $N_k$ sub-time steps have been completed. If the C cell does not divide, it moves according to the rules described in the section 2.3.2.

When the $N_k$ sub-time steps have been repeated independently, we add to the initial population cells that are either born by proliferation or recruited by infiltration. Dead cells are removed. The number of cells is then $N_{k+1}$. We start a new cycle of $N_{k+1}$ sub-time steps, as previously described. Therefore, over a time step, a given cell will on average die, move or divide (if it is a C cell).

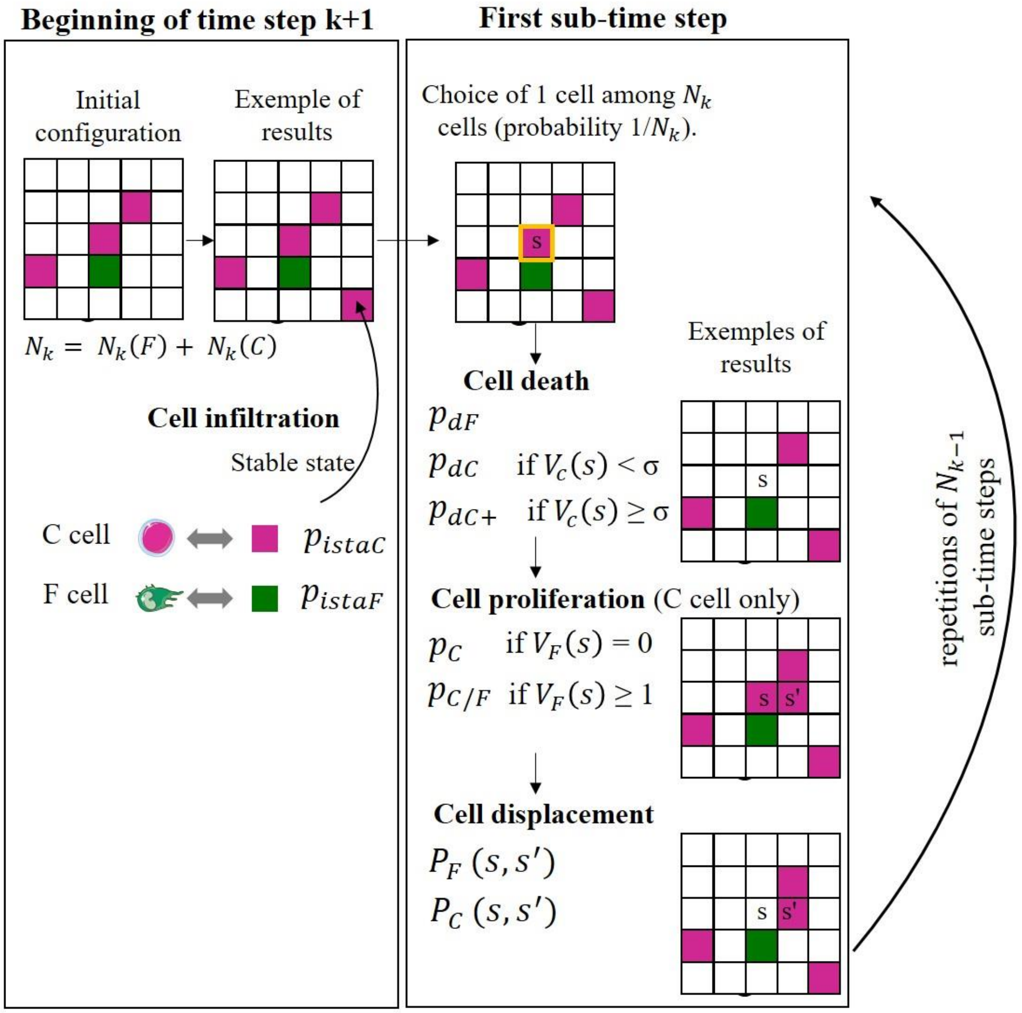


Fig. F. Examples of cell infiltration, death, proliferation and displacement of cells at each time step. We consider the beginning of the time step $k + 1$. This period is divided into $N_k$ sub-time steps, where $N_k$ is the number of cells at the beginning of period $k + 1$. F and C cells are indicated by respectively green and purple squares. Adapted from (Dupin et al., 2023).

Infiltration during exacerbations:

Every year, i.e. after 175,200 time steps, we add a number $N_{iexaF}$ of F cells with the probability $p_{iexaF}$. If F cells are recruited by infiltration during an exacerbation, they are randomly and uniformly positioned among all of the vacant sites. Cells that have infiltrated in the meantime are added to the initial population. A new time step starts again as described above.

### 3 Determination of parameters using biological informations

For a control subject as for a patient with COPD, the same model is ~~be~~ applied but with different parameters. These parameters are estimated thanks to experiments and data from the literature and are defined below (see Table S11 for numerical values). Simulations are performed using these parameter values.

For the estimation of $v_0$, $p_{dC}$, $p_{dF}$, $x_C$, $x_F$, $p_{istaC}$ $p_{istaF}$, $p_{iexaF}$ and $N_{iexaF}$, a streamlined model has been used, in which a mathematical and probabilistic analysis was possible (Dupin et al., 2023). The major interest of the streamlined model is that it allows a rigourous mathematical analysis to accurately calculate parameters. We note $\delta^{Ctl}$ and $\delta^{COPD}$ the value of the parameter $\delta$ for respectively a control subject and a patient with COPD.

The streamlined model is a particular case of the model, where local interactions, *i.e* C cell-induced cell death and contact inhibition of C cell proliferation play no role. To reflect these two properties, σ and λ have been fixed respectively to 9 and 0. This means that C and F cell displacement, the infiltration at the stable state and during exacerbations are identical to the complete model.

#### *3.1 Determination of the cell death parameters*

##### 3.1.1 Determination of $p_{dF}$

Due to the inability to easily label living F cells *in vivo*, there is no quantitative data on the lifespan of these cells. We will therefore approximate that F cells, as cells derived from monocytes, have a lifespan $hl(F)$ similar to those of interstitial macrophages (derived from monocytes) in the lungs. A recent study shows that the half-life of interstitial macrophages derived from monocytes is approximately 10 months in a healthy context (Schyns et al., 2019).

For a control subject, we choose the value of the probability $p_{dF}^{Ctl}$ of dying for an F cell over a period of 3 min equal to those calculated in the streamlined model using the $hl^{Ctl}(F)$ lifespan of a F cell in the control case (Dupin et al., 2023):

$$p_{dF}^{Ctl} = 1 - e^{-ln2/hl^{Ctl}(F)} \approx 4.8\times10^{-6}$$

Our previous work showed that the exposure of fibrocytes to the secretions of the bronchial epithelia of patients with COPD decreases by a factor 2 the percentage of dead cells (Dupin et al., 2019). In patients with COPD, we will therefore choose for each F cell the probability $p_{dF}^{COPD}$ of dying equal to :

$$p_{dF}^{COPD} = p_{dF}^{Ctl}/2 \approx 2.4\times10^{-6}$$

#### 3.1.2 Determination of $p_{dC}$

Data obtained in murine lungs show that CD8$^{+}$ cells have a half-life $hl(C)$ of 14 days in the lung (McMaster et al., 2015). For a control subject, we choose the value of the probability $p_{dC}^{Ctl}$ of dying for an C cell over a period of 3 min equal to those calculated in the streamlined model using the $hl^{Ctl}(C)$ lifespan of a C cell in the control case (Dupin et al., 2023):

$$p_{dC}^{Ctl} = 1 - e^{-ln2/hl^{Ctl}(C)} \approx 1.0\times10^{-4}$$

A study suggests a modification of the C cell death processes in tissues from patients with COPD, with decrease by about half of the percentage of apoptotic C cells in the distal airways of mild to moderate COPD patients (Siena et al., 2011), which constitute the majority of our study cohort. In patients with COPD, we will therefore choose for each C cell the probability $p_{dC}^{COPD}$ of dying equal to :

$$p_{dC}^{COPD} = p_{dC}^{Ctl}/2 \approx 5\times10^{-5}$$

#### 3.1.3 Determination of $p_{dC+}$ and σ

Our image analyzes on lung tissues showed that clusters of C cells contain a median value of approximately 4 cells (Figure 1). Thus, we choose σ = 3 neighbors, to favor cell death for a neighbourhood comprising 3 or more C neighbors.

A recent study demonstrated the existence of a negative feedback loop acting by CTLA-4, to limit the expansion of activated lymphocytes (Zenke et al., 2020). Cross-linking of CTLA-4 to the surface of lymphocytes induces apoptosis of previously activated lymphocytes, going from less than 5% of apoptotic lymphocytes to about 90% (Gribben et al., 1995). These quantifications were refined in another study, which shows that the cross-linking of CTLA-4 induces an increase in the number of apoptotic lymphocytes of about 4 times (Scheipers & Reiser, 1998). We therefore choose:

$$p_{dC+} = 4 \times p_{dC}$$

The increased probability $p_{dC+}$ of dying when a C cell has many other C cells in its neighbourhood is equal to 4 times the probability $p_{dC}$, regardless of the condition. We thus obtain :

$$p_{dC+}^{Ctl} = p_{dC}^{Ctl} \times 4 \approx 4.0 \times 10^{-4}$$

$$p_{dC+}^{COPD} = p_{dC}^{COPD} \times 4 \approx 2.0 \times 10^{-4}$$

Our analyzes on lung tissues showed that clusters of C cells contain a median value of approximately 4 cells, in patients with COPD as in control subjects (Figure 1), suggesting that σ is unchanged in patients with COPD vs control subjects :

$$\sigma^{Ctl} = \sigma^{COPD} = 3$$

*3.2 Determination of the cell proliferation parameters*

#### 3.2.1 Determination of $p_F$

Based on our own unpublished observations and published studies (Ling et al., 2019; Schmidt et al., 2003), fibrocytes very poorly proliferate in culture, allowing us to consider that an F cell does not divide in lung tissue. We will consider that the probability $p_F$ of dividing for an F cell over a period of 3 min is identical in control subjects and COPD patients and equal to :

$$p_F^{Ctl} = p_F^{COPD} = 0$$

#### 3.2.2 Determination of $p_C$

The cell cycle length of a C cell is highly variable, and it is difficult to obtain quantitative data at steady state. During the 3 to 6 weeks following an immunization, lung C cells undergo 1 to 2 cycles of proliferation (Bivas-Benita et al., 2013). We will then consider an average duration of 6 weeks = 6 × 7 × 24 × 60 = 60 480 min = 20 160 × 3 min for a cell cycle. We obtain the probability $p_C$ that a C cell divide during the 3 min-time step, for control subjects as well as for patients with COPD :

$$p_C^{Ctl} = p_C^{COPD} = 1/20\ 160 \approx 5.0 \times 10^{-5}$$

#### 3.2.3 Determination of $p_{C/F}$

Our *in vitro* experiments show that the presence of fibrocytes multiplies the rate of C cells, which have been divided, by approximately 4, if we consider respectively C cells previously activated by beads coated with anti-CD3 and anti-CD28 antibodies. The increased probability $p_{C/F}$ of dividing will therefore be taken equal to 4 times the value of the probability $p_C$. This probability is identical in control subjects and patients with COPD :

$$p_{C/F}^{Ctl} = p_{C/F}^{COPD} = 1/5040 \approx 2.0 \times 10^{-4}$$

*3.3 Determination of cell displacement parameters*

3.3.1 Determination of $f_F$

Our chemotaxis experiments show that F cells are almost not attracted towards the secretion of C cells purified from control subjects (Figure 2). This justifies an almost zero attraction ($\varepsilon_F$, « small », arbitrarily chosen as $\varepsilon_F$ $=10^{-3}$).

For control subjects, we have thus choosen for $f_F^{Ctl}(n)$ (Fig. G):

$$f_F^{Ctl}(n) = \varepsilon_F, \qquad if\ n \in \{0, 1, 2, 3, 4\ 5, 6, 7, 8\}$$

In the COPD situation, our experiments also show that F cells are significantly attracted towards the secretion of C cells obtained from patients with COPD (Figure 2). As this chemotactic effect requires soluble factors that have to secreted in a sufficient concentrations, this justifies an almost zero attraction for s' such as $V(C)(s') < 4$ cells and a maximal and constant attraction for s' such as $V(C)(s')$= 4 or 5 cells. On the other hand, the attraction of the site s' for a F cell probably decreases when the site is too "crowded", because of physical hindrance and/or or the secretion of factors that are secreted when many C cells are aggregated. This is in agreement with the theory of quorum sensing (Antonioli et al., 2018) and with our observations that the median number of cells in clusters containing F and C cells is relatively low (6 for patients with COPD). This leads us to choose an almost zero attraction for s' such as $V(C)(s') > 5$ cells. Thus, a F cell will preferentially and uniformly go to an empty s' site such that $V(C)(s')$ = 4 or 5, or will stay on the site s.

For patients with COPD, we have thus chosen for $f_F^{COPD}(n)$ (Fig. G):

$$f_F^{COPD}(n) = \begin{cases} 1, & if\ n = 4\ or\ 5 \\ \varepsilon_F, & if\ n \in \{0, 1, 2, 3, 6, 7, 8\} \end{cases}$$

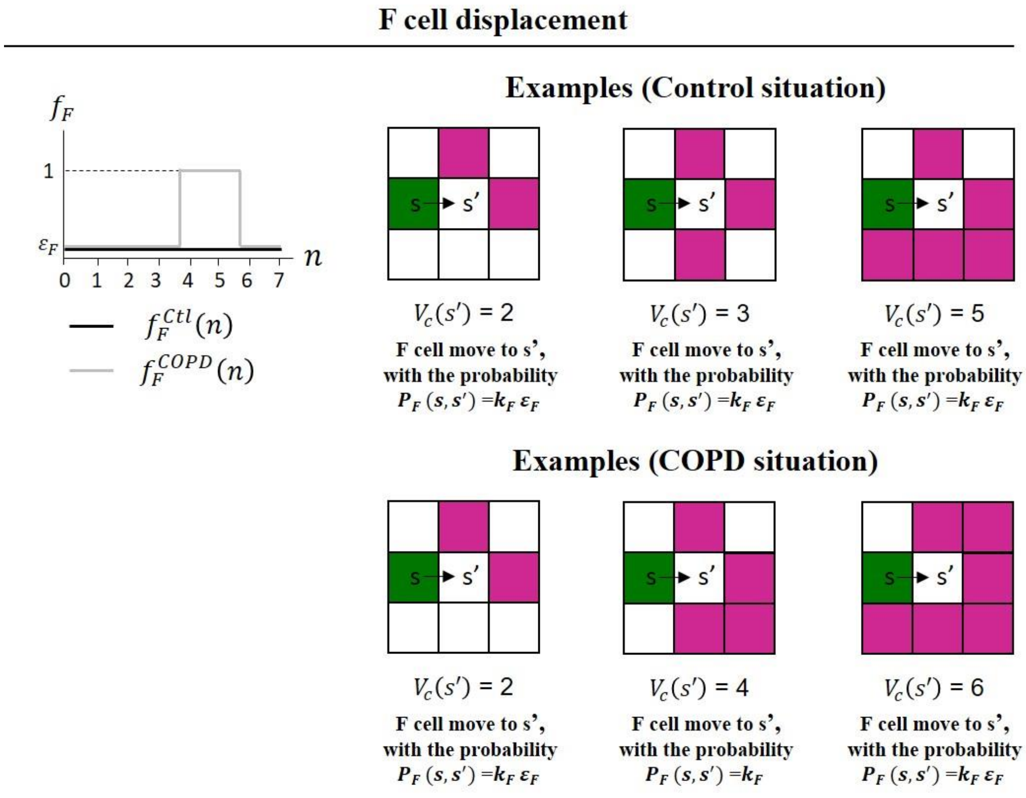


Fig. G. F Cell displacement rules. F and C cells are indicated by respectively green and pink squares. $\varepsilon_F$ has been taken equal to $10^{-3}$

#### 3.3.2 Determination of $f_C$

For control subjects, we have thus chosen for $f_C^{Ctl}(n)$ :

$$f_C^{Ctl}(n) = \begin{cases} 1, & if\ n = 4\ or\ 5 \\ \varepsilon_C, & if\ n\ \in \{1,2,3,6,7,8\} \end{cases}$$

with $\varepsilon_C = 10^{-3}$

Based on the same type of justifications than those used for $f_F$ , the choice of $f_C$ reflects an almost zero attraction for s' such as $V(s')$<4 cells or $V(s')$>5 cells and a maximal and constant attraction for s' such as $V(s')$=4 or 5 cells. As $V(s')$ includes the C cell considered at the site s and is therefore $\geq 1$, $f_C$ is defined for n $\in \{1,2,3,4,5,6,7,8\}$.

We will consider that $f_C$ is identical in control subjects and COPD patients : $f_C^{Ctl}(n) = f_C^{COPD}(n)$

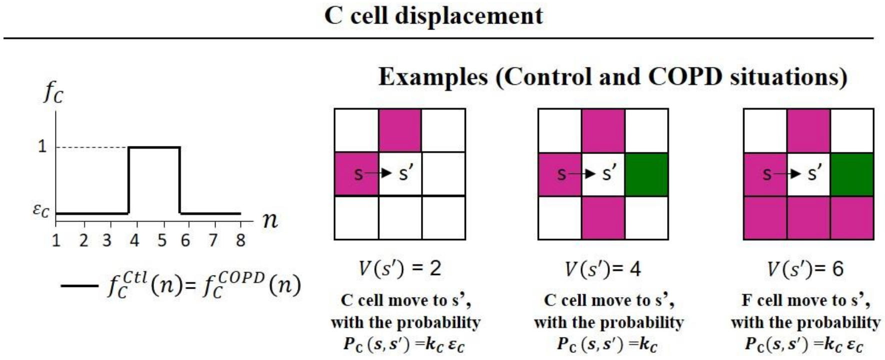


Fig. H. C Cell displacement rules. F and C cells are indicated by respectively green and pink squares. $\varepsilon_F$ has been taken equal to $10^{-3}$. Adapted from (Dupin et al., 2023).

#### 3.3.3 Determination of $x_F$ and $x_C$

The parameters $x_F$ and $x_C$ are terms involved in the definition of respectively $P_F\,(s,s')$ and $P_C\,(s,s')$. They are chosen such as the median speed of F and C cells is equal to 2.3 µm/min. We have previously demonstrated using the streamlined model (Dupin et al., 2023) that :

$$x_F = \frac{1-\alpha/5}{\alpha/5}\left(\sum_{s'\in M(s)} f_F\left(V(C)(s')\right)1_{\{s'empty\}}\right),$$

with $\alpha$ being a real number, whose numerical value is $\alpha = 4.67091$.

This implies that:

$$k_F = \frac{\alpha/5}{\sum_{s'\in M(s)} f_F\left(V(C)(s')\right)1_{\{s'empty\}}}$$

For the C cells, the real numbers $x_C$ and $k_C$ are obtained similarly :

$$x_C = \frac{1-\alpha/5}{\alpha/5}\left(\sum_{s'\in M(s)} f_C\left(V(s')\right)1_{\{s'empty\}}\right),\ k_C = \frac{\alpha/5}{\sum_{s'\in M(s)} f_C\left(V(s')\right)1_{\{s'empty\}}}$$

These parameters are identical for control subjects and patients with COPD.

*3.4 Determination of cell infiltration parameters*

3.4.1 Determination of $p_{istaF}$, $p_{iexaF}$ and $N_{iexaF}$

F cells have a limited lifespan in the lungs, with a half-life that we have estimated at 10 months in the lung, by analogy with the half-life of interstitial macrophages (Schyns et al., 2019). Our previous work indicates presence of F cells in the lungs, at varying densities in control subjects and COPD patients (Dupin et al., 2019), suggesting infiltration of F cells at stable state, in order to maintain this pulmonary pool relatively constant. This leads us to add, for a healthy subject as for a COPD subject, F cells with a probability $p_{istaF}$.

We have demonstrated in the streamlined model (Dupin et al., 2023) that the expectation of $N_k(F)$ is close to $N_0(F)$ for large $k$ is realized if $p_{istaF}$ = $p_{dF}$ $N_0(F)$. We kept this relation in the full model for the control situation. With $N_0(F) = 19$, and $p_{dF}^{Ctl} \approx 4{,}8\times10^{-6}$, we obtain for a control subject:

$$p_{istaF}^{Ctl} \approx 9.12\times10^{-5}$$

The level of circulating fibrocytes is identical in control subjects and patients with COPD at the stable state (Dupin et al., 2016), suggestion that $p_{istaF}$ is identical in control subjects and COPD patients :

$$p_{istaF}^{COPD} = p_{istaF}^{Ctl} \approx 9.12\times10^{-5}$$

The choice of $p_{istaC}$ is different, see section 3.4.2.

The value of $p_{iexaF}$ depends closely on the condition of the subject (healthy *vs* COPD). For a healthy subject, as there is no exacerbation, this probability is zero :

$$p_{iexaF}^{Ctl} = 0$$

For COPD patients, this value is non-zero. Indeed, we previously showed that there is an increase in the concentration of F cell level in peripheral blood of COPD patients during acute exacerbation (Dupin et al., 2016). In addition, in the lungs, the density of F cells is higher in the tissues of COPD patients than in those of healthy subjects (Dupin et al., 2019). Since F cells proliferate little or not at all (our own unpublished observations and published studies (Ling et al., 2019; Schmidt et al., 2003), this suggests that for COPD patients, F cells are recruited from the blood to the lungs at the time of exacerbations. To take this into account, for COPD patients, we will add a number $N_{iexaF}$ of F cells in the surface of interest with the probability $p_{iexaF}$ at a frequency corresponding to the average exacerbation frequency of COPD patients, so that after 20 years, on average, the number of F cells is double in COPD patients than in healthy patients. The average number of exacerbations is 0.85, 1.34 and 2/year in patients with stages 2, 3 and 4 COPD, respectively (Hurst et al., 2010). In our cohort for tissue analysis, the majority of patients are stage 2 or 3, so we choose an average number of exacerbations of 1 per year. We will achieve the goal of doubling F cells after 20 years by adding enough F cells each year to compensate for cell death. We agree that the first addition takes place in year 0 and the last is done in the nineteenth year. We do the same assumption as for infiltration at stable state: if we add $N_{iexaF}$ at the start of a year, these cells are not active immediately, it is necessary to wait for the start of the following time step for them to be.

The goal is to choose $p_{iexaF}^{COPD}$ in such a way that the expectation of $N_T(F)$ is the double of $N_0(F)$, where T = 20 years = 3 504 000 time steps. Let us remind the important fact that the F cells die independently of the environment and do not duplicate themselves. Therefore we can realize the condition of doubling the number of F cells after20 years using the streamlined model.

We then get (Dupin et al., 2023):

$$N_{iexaF} = \left[\frac{K_1}{K_2}\right] + 1 \quad \text{and} \quad p_{iexaF} = \frac{1}{N_{iexaF}}\frac{K_1}{K_2}$$

With:

$$K_1 = (1-(1-p_{dF}^{COPD})^{a1})\left(2N_0(F) - (1-p_{dF}^{COPD})^T\left(N_0(F) - \frac{p_{istaF}^{COPD}}{p_{dF}^{COPD}}\right) - \frac{p_{istaF}^{COPD}}{p_{dF}^{COPD}}\right)$$

$$K_2 = (1-p_{dF}^{COPD})^{a1-1}(1-(1-p_{dF}^{COPD})^T)$$

And

$$a1 = 1\ year = 175\ 200\ time\ steps\ of\ 3\ min$$

With the values of the previously determined parameters we obtain:

$$N_{iexaF}^{COPD} = 1, p_{iexaF}^{COPD} \approx 0.0022$$

#### 3.4.2 Détermination of $p_{istaC}, p_{iexaC}$ and $N_{iexaC}$

Since C cells have a limited lifespan in the airways, with an estimated half-life of 14 days in the lung (McMaster et al., 2015), it has been proposed that the number of memory C cells in the lung tissue is maintained through continuous recruitment either from the systemic pool of cells (Ely et al., 2006) or from niches in the pulmonary interstitium (Takamura et al., 2016), rather than by proliferation or prolonged survival. This leads us to add C cells by infiltration in such a way that the number of C cells fluctuate little (at least at the equilibrium). The estimation of the probability of infiltration of C cells at steady state $p_{istaC}$ is described below. We first consider the control situation.

Recall that in the streamlined model (2) if we choose $p_{istaC} = p_{dC} \times N_0(C)$ then the mean number of cells C remains constant at equilibrium. According to the values of $p_{dC}^{Ctl}$, we have $p_{dC}^{Ctl}\ N_0(C) \approx 1.18\ \times 10^{-2}$

We realize $K$ independent simulations of our model up to the time step $T+1$. We are interested in the $N_T$ sub-steps which follow the first $T$ steps.

For each simulation $1 \leq i \leq K,\ N_{T,i}$ stands for the number of cells at time $T$.

For any $0 \leq\ l\ \leq\ N_{T,i}$, let us introduce:

$\alpha^{-}_{l,T,i}$ = the number of $C_{\mathrm{j}} -$ cells, computed at the end of the sub-step l, which are alive, did not duplicate and $V(C_j) < \sigma$,
$\alpha^{+}_{l,T,i}$ = the number of $C_{\mathrm{j}} -$ cells, computed at the end of the sub-step l, which are alive, did not duplicate and $V(C_j) \geq \sigma$,
$\beta_{l,T,i}$ = the number of $C_{\mathrm{j}} -$ cells, computed at the end of the sub-step l, which are alive, did not duplicate, there exists $\mathrm{s}' \in \mathrm{M}(C_j)$, $\mathrm{s}'$ empty, and $V(C)(s') < \lambda$ and there exists no $F$ in $\mathrm{M}(C_j)$,
$\beta^{+}_{l,T,i}$ = the number of $C_{\mathrm{j}} -$ cells, computed at the end of the sub-step l, which are alive, did not duplicate, there exists $\mathrm{s}' \in \mathrm{M}(C_j)$, $\mathrm{s}'$ empty, and $V(C)(s') < \lambda$ and there exists at least one $F \in \mathrm{M}(C_j)$.
and

$$A^{+}_{T} = \frac{1}{K}\left\{\sum_{i=1}^{K} \frac{1}{N_{T,i}}\left(\sum_{l=0}^{N_{T,i}-1} \alpha^{+}_{l,T,i}\right)\right\} A^{-}_{T} = \frac{1}{K}\left\{\sum_{i=1}^{K} \frac{1}{N_{T,i}}\left(\sum_{l=0}^{N_{T,i}-1} \alpha^{-}_{l,T,i}\right)\right\}$$

$$B_{T} = \frac{1}{K}\left\{\sum_{i=1}^{K} \frac{1}{N_{T,i}}\left(\sum_{l=0}^{N_{T,i}-1} \beta_{l,T,i}\right)\right\} B^{+}_{T} = \frac{1}{K}\left\{\sum_{i=1}^{K} \frac{1}{N_{T,i}}\left(\sum_{l=0}^{N_{T,i}-1} \beta^{+}_{l,T,i}\right)\right\}$$

We agree that sub-step $0$ is at the end of the period $T$.

Under these conditions, we can prove:

$$\mathbb{E}(N_{k+1}(C)) \approx \mathbb{E}(N_k(C)) + p_{istacC} + p_C B_T + p_{C/F} B^{+}_{T} - p_{dC^+} A^{+}_{T} - p_{dC} A^{-}_{T}. \qquad (1)$$

For any period $k$, where $\mathbb{E}(N_k(C)$ and $\mathbb{E}(N_{k+1}(C))$ is the mean number of C cells at the time step k (resp. k+1).

**Remark**

The coefficients $A^{+}_{T}$, $A^{-}_{T}$, $B_T$ and $B^{+}_{T}$ can be interpreted as follows: during the $N_T$ sub-steps following the period $T$,

- $A^{+}_{T}$ (resp. $A^{-}_{T}$) is the average number of times C cells can potentially die with probability $p_{dC+}$ (resp. $p_{dC}$)

- $B_T$ (resp. $B_T^+$) is the average number of times C cells can potentially proliferate with probability $p_C$ (resp. $p_{C/F}$).

All the parameters are given by experimental data, the only one which is free is $p_{istaC}$. However, the relation (1) does not permit to determine $p_{istaC}$, because $B_T$, $B_T^+$, $A_T^+$ and $A_T^-$ also depend on $p_{istaC}$. Our strategy is the following: we fix $T$ and we we want to determine $p_{istaC} = p_{istaC}(T)$ such that

$$\mathbb{E}(N_{T+1}(C)) = \mathbb{E}(N_T(C)) \tag{2}$$

From (1) this condition is equivalent to:

$$p_{istaC}(T) = p_{dC+} \times A_T^+ + p_{dC} \times A_T^- - p_C \times B_T - p_{C/F} \times B_T^+. \tag{3}$$

We read (3) as an equation of the form:

$$p_{istaC}(T) = \phi(p_{istaC}(T)) \tag{4}$$

where $\phi$ is a function and we solve it by iteration. We begin with $p_{istaC}^0(T)$ as the value of $p_{istaC}^{ctrl}$ in the streamlined model (Dupin et al., 2023), $p_{istaC}^0(T) = 1.18 \times 10^{-2}$.

We simulate $K$ times the complete model over the time period $[0, T+1]$ and we get:

$$p_{istaC}^1(T) = \phi(p_{istaC}^0(T)). \tag{6}$$

We repeat this process, then we define by induction a sequence $(p_{istaC}^k(T))_k$ as

$$p_{istaC}^{k+1}(T) = \phi(p_{istaC}^k(T)). \tag{7}$$

We stop the iterations as soon as $k \mapsto p_{istaC}^k$ "seems" constant. Let $p_{istaC}^{ctrl}$ be this value.

Beware that the condition (2) does not imply that $\mathbb{E}(N_{T'}(C)) = \mathbb{E}(N_T(C))$ for $T' > T$. We verify by simulations that

$$\mathbb{E}(N_{T'}(C)) \approx \mathbb{E}(N_T(C)), \quad for\ any\ \ T \leq T' \leq T_{20y} \tag{8}$$

where $T_{20y} = 20\ \ years\ \ = 3\ \ 504\ \ 000$ time steps.

As $T_{20y}$ represents a large number of time steps we will aggregate them month by month. Let $T_m$ be the number of periods to obtain one month:

$$T_m = \frac{T_{20y}}{240} = 146\ \ 000\ \ periods\,.$$

We simulate $K$ times our model (with parameter $p_{istaC}^{ctrl}$ determined as explained above) up to time $T_{20y}$. First, we consider the stochastic fluctuations in the number of $C$ cells. For any simulation $1 \leq i \leq K$ and month $0 \leq j < 240$, let

$$\overline{N}_{jT_m,i}(C) = \frac{1}{T_m}(\sum_{l=0}^{T_m-1} N_{jT_m+l,i}(C))$$

where $N_{k,i}(C)$ is the number of $C$ cells at the end of the $k$ time steps, and for the simulation $i$.

It is also interesting to compare the family of curves $((\overline{N}_{jT_m,i}(C))_{0\leq j<240}, 1 \leq i \leq K)$ with the variations of the mean values $(\overline{N}_{jT_m}(C))_{0\leq j<240}$ where:

$$\overline{N}_{jT_m}(C) = \frac{1}{K}(\sum_{i=1}^{K} \overline{N}_{jT_m,i}(C\text{to}\ \overline{N}_T(C))), \quad 0 \leq j < 240.$$

We observed that numerically $\overline{N}_T(C)$ is close to $\overline{N}_{T_{20y}}(C)$.

Of course, it remains to choose a value for T. In all the simulations, we found numerically that the stationary state was reached rather quickly, by following the fluctuations of the numbers of cells C and F. We have chosen T = 2 years.

Finally, we obtain the following estimation for $p_{istaC}^{Ctl}$ :

$$p_{istaC}^{Ctl} = p_{dC}^{Ctl}\ n_0(C) \approx 1.40\times10^{-2}$$

There is no biological evidence of difference in the level of circulating C cells between healthy subjects and patients with COPD, suggestion that $p_{istaC}$ is identical in control subjects and COPD patients :

$$p_{istaC}^{Ctl} = p_{istaC}^{COPD} \approx 1.40\times10^{-2}$$

Concerning $p_{iexaC}$, the literature shows that there is probably an infiltration of C cells in the lungs, especially in COPD patients (Freeman et al., 2007; Saetta et al., 1999), but the relationships with exacerbations are not entirely clear. For simplification, we will assume that there is no C cell infiltration during exacerbations. Thus, for healthy subjects as well as for patients with COPD the value of $p_{iexaC}$ is zero :

$$p_{iexaC}^{Ctl} = p_{iexaC}^{COPD} = 0$$

#### *3.5 Determination of parameters upon therapeutic interventions*

##### 3.5.1 Determination of parameters upon CXCR1/2 inhibition

Our results show reparixin (dual CXCR1/CXCR2 antagonist) treatment completely suppress the increased chemotaxis induced by the secretions of $CD8^+$ T cells purified from COPD lungs (Figure 2E). We will thus choose:

$$f_F^{CXCR1/2\ inhibition}\ (n) = f_F^{Ctl}(n) = \varepsilon_F\,, \qquad if\ n\ \in\ \{0,1,2,3,4\ 5,6,7,8\}$$

All the others parameters remain unchanged and similar to COPD situation.

##### 3.5.2 Determination of parameters upon CD86 or CD54 inhibition

Our results show the treatment with CD54 and CD86-blocking antibodies in the co-culture assay significantly reduced the fibrocyte-induced proliferation of $CD8^+$ T cells by a factor respectively 1.5 and 1.2 (Figure 4).

We will thus choose:

$p_{C/F}^{CD54\ inhibition} = p_{C/F}^{COPD}/1{,}5 = 1/(5040 \times 1.5) = \approx 1.3 \times 10^{-4}$

$p_{C/F}^{CD86\ inhibition} = p_{C/F}^{COPD}/1{,}2 = 1/(5040 \times 1.2) = \approx 1.6 \times 10^{-4}$

All the others parameters remain unchanged and similar to COPD situation.

### 4 Simulation procedure

Our algorithm (see section 2.6 and Fig. F) is implemented in Julia, which allows to parallelize the computation sequences as well as to use graphic libraries, in order to produce drawings and videos illustrating step by step the evolution of a starting situation according to the biological parameters selected to launch the simulations. Our program is modular, in the sense that it is made up of reusable functions for the benefit of users wishing to test other configurations involving different evolutionary laws. A complete version of the program can be downloaded from the following site: https://plmbox.math.cnrs.fr/d/49bcbc1db63a4654be7e/

### 5 Definition of measurements at the final state

#### *5.1 Quantification of the density of C cells in interaction with F cells*

For each F cell in a site $(i, i) \in P$, the number of C cells belonging to the the set of $(i-1, j), (i, j-1), (i, j+1), (i+1, j)$ is counted. The density of C cells in interaction with F cells corresponds to the sum of these numbers of neighbouring C-cells divided by the area of the lamina propria (0.179 $mm^2$).

#### *5.2 Quantification of the minimal distances between C cells and F cells*

For each F cell in a site $= (i, j) \in$ P, we set $M^{(0)}(s) = M(s)$ and for any integer k $\geq$ 1:

$$M^{(k)}(s) = \{(i', j') \in P, max\ |i\ -\ i'|, |j - j'|\ \}\ \leq\ k\}$$

Obviously $M^{(0)}(s)\ \subset\ M^{(1)}(s)\ \subset \cdots\ \subset\ M^{(k)}(s)$. $M^{(k+1)}(s)$ is obtained by adding another layer of lattice sites around $M^{(k)}(s)$.

Consider a C cell located in s'. We define as the minimal distance between the considered F cell and the C cell the number: $k\ \times 7$µm if s $\in M^{(k)}(s)$ and s $\notin\ M^{(k+1)}(s)$

The same process is repeated until a C cell is founded. For each F, there a minimal distance associated. For each simulation at the final state, the mean value of the minimal distances can be calculated, and the frequency distributions of minimal distances can also be determined.

#### *5.3 Quantification of the clusters*

Clusters of cells are defined for a number of cells greater or equal to 2. Cells belong to the same cluster if they are in contact by a corner or a side. Clusters are automatically recorded at each time step and at the final state of the simulations. The distinction between clusters containing exclusively C cells, F cells, or both cell type (“mixed” clusters) is also done. The number of cell in each cluster is also recorded. The mean number of cells per cluster is the average of the number of cells for all the clusters detected at the considered time step. The density of clusters corresponds to the number of clusters divided by the area of the lamina propria (0.179 $mm^2$).

### Supplementary figures

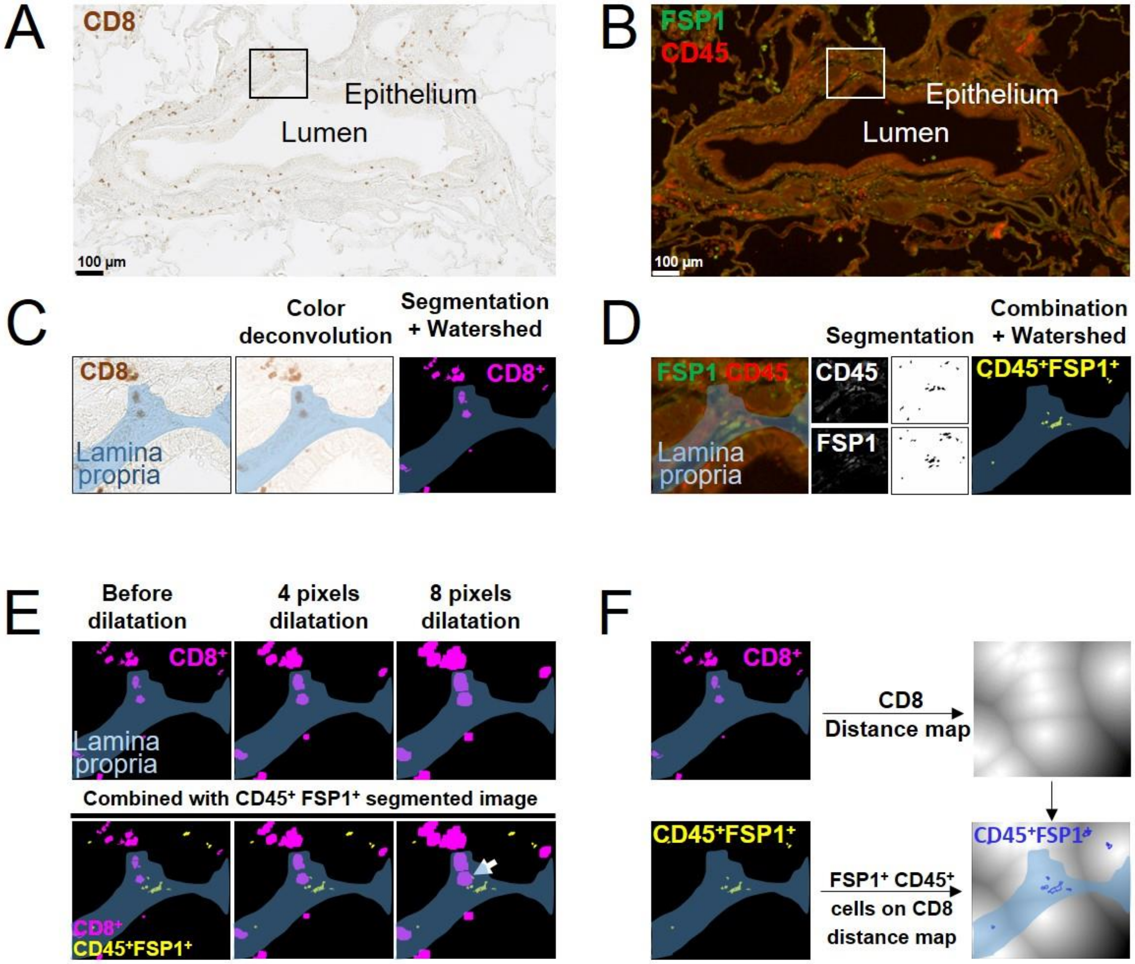


**Figure S1. Detection of CD8$^{+}$ T cells, CD45$^{+}$ FSP1$^{+}$ cells, and quantification of interacting cells, and the minimal distances between the two cell types.** (**A**) CD8 (brown) staining of a representative bronchus. (**B**) CD45 (red) and fibroblast-specific protein 1 (FSP1, green) stainings of the same bronchus. (**C, D**) The left panels show a higher magnification image (indicated by the boxed regions) of the images in **A** and **B**. The lamina propria is shown in light blue. (**C**) Middle panel: image for CD8 staining obtained after colour deconvolution. Right panel: image obtained after segmentation by a binary threshold followed by a watershed transformation to the segmented image. CD8$^{+}$ T cells are shown in magenta. (**D**) Middle panels: images for CD45 (top) and FSP1 (bottom) stainings and images obtained after segmentation by a binary threshold. Right panel: segmented image obtained after combination of CD45 and FSP1 segmented images to select cells dual positive for FSP1 and CD45 double staining, followed by a watershed transformation to separate potential neighboring cells. CD45$^{+}$ FSP1$^{+}$ cells are shown in yellow. (**E**) Top panels: dilatation of each CD8 positive particle with an area greater than 64 µm$^{2}$. Bottom panels: combination of these modified images with segmented image for dual CD45 FSP1 positive staining (**B**) to automatically select overlapping dilated CD8$^{+}$ T cells with CD45$^{+}$ FSP1$^{+}$ cells. The white arrow indicates interacting cells. (**F**) Top panels: a CD8 distance map is built from the binary image produced from CD8 staining. Bottom panels: each area corresponding to a FSP1$^{+}$ CD45$^{+}$

cell was reported on the CD8 distance map (blue outlines), and the minimal gray value in each area is measured and converted to a distance, allowing to measure the minimal distance between the $CD45^+$ $FSP1^+$ cell and neighbouring $CD8^+$ T cells.

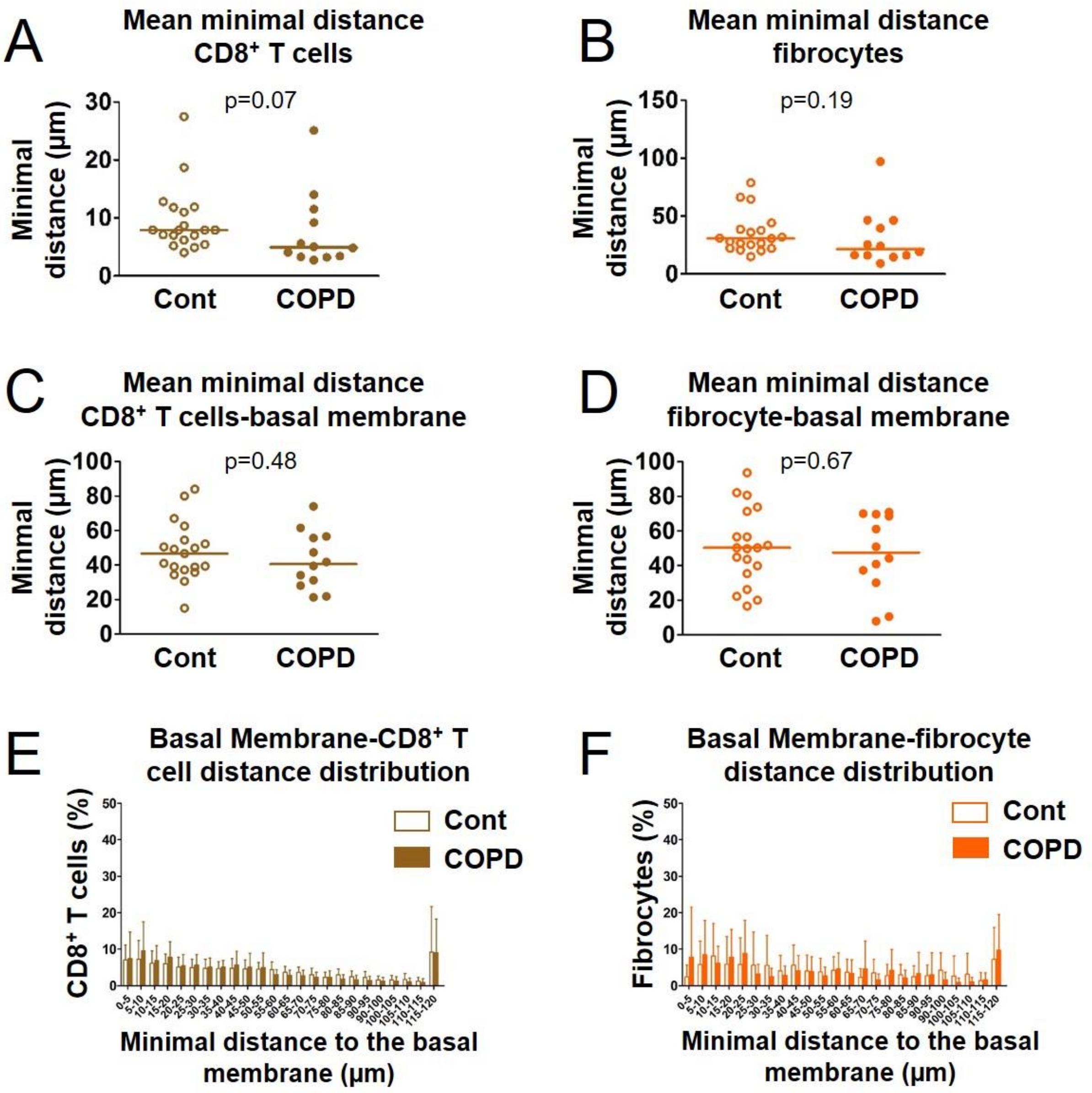


**Figure S2. Spatial distributions of CD8$^+$ T cells and fibrocytes in the peribronchial area.** (**A**, **B**) Quantification of the mean minimal distances between CD8$^+$ T cells (A) and between fibrocytes (**B**) in one specimen/patient. (**C**, **D**) Quantification of the mean minimal distances between CD8$^+$ T cells and the basal membrane (**C**), and between fibrocytes and the basal membrane (**D**) in one specimen/patient. (**A**-**D**) The medians are represented as horizontal lines, n=20 specimens from control subjects, n=12 specimens from patients with COPD. Mann Whitney tests. (**E**, **F**) Mean frequency distribution of minimal distances (with 5 µm binning) between CD8$^+$ T cells and the basal membrane (**E**) and between fibrocytes and the basal membrane (**F**) for control subjects (white) and COPD patients (brown/orange). Error bars indicate standard error of the mean. Two-way ANOVA for repeated measures: $F(23, 696)=0.48$, $P=0.98$ (**E**), $F(23, 696)=1.10$, $P=0.34$ (**F**).

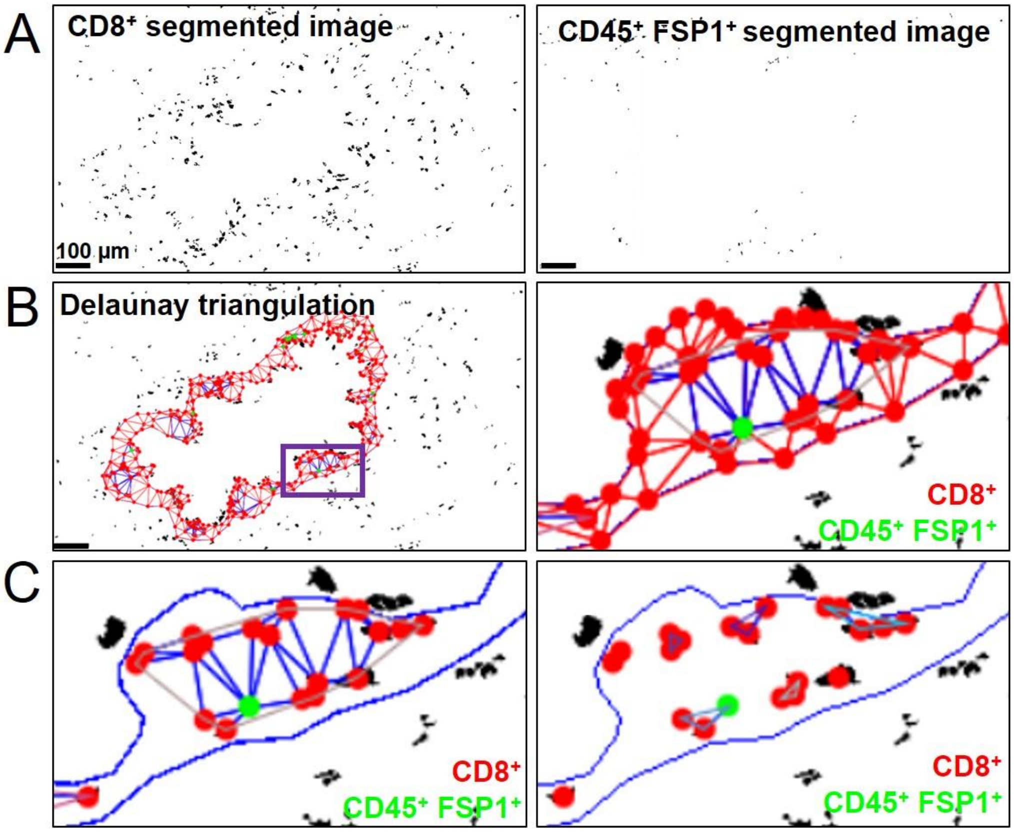


**Figure S3. Principle of the use of Delaunay triangulation for cluster analysis. (A**) Binary images for CD8 (left panel) and double CD45-FSP1 (right panel) stainings obtained after segmentation. (**B**) Images after Delaunay triangulation, performed on the centers of mass of CD8$^+$ T cells and fibrocytes and on the points defining area edges. CD8$^+$ T cells and fibrocytes appear respectively with red and green dots. The right panel is a higher magnification of the peribronchial area (indicated by the boxed purple region on the left panel). The connections including the points defining area edges are shown in red, all the other connections are shown in blue. (**C**) Left panel: image with Delaunay triangulation after elimination of the connections including the points defining area edges. The lamina propria is shown in blue. Right panel: image shown on the left panel, after applying a threshold value (40 µm) above which connections are not kept.

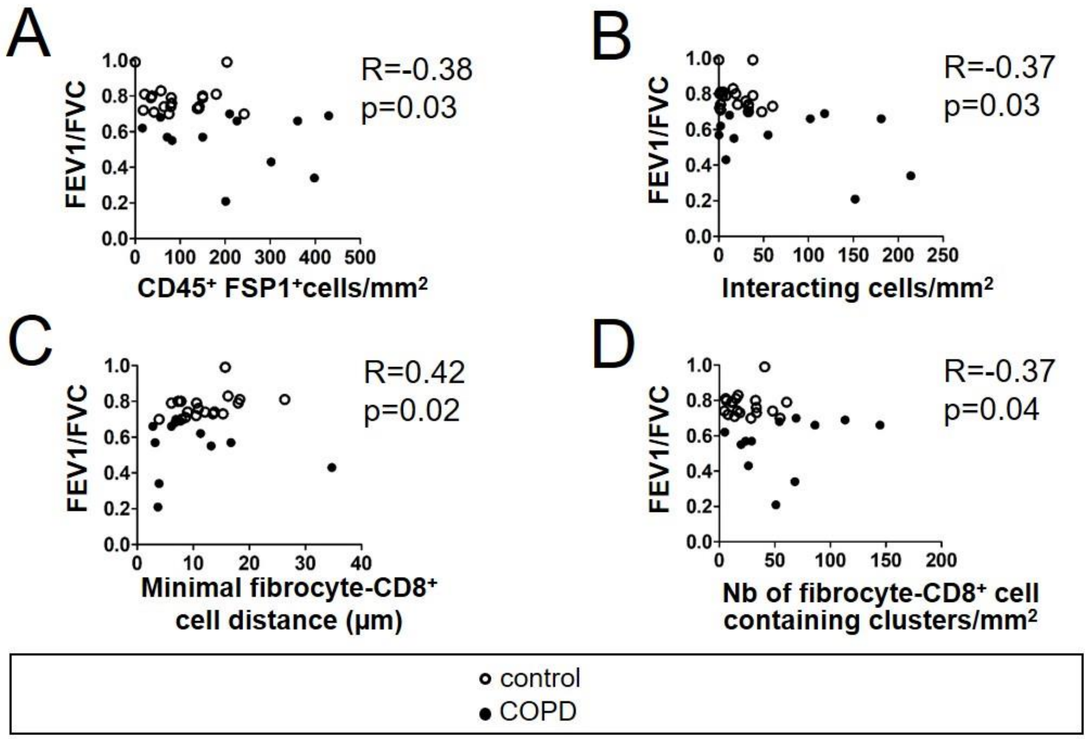


**Figure S4. Relationships between the $FEV_1$/FVC ratio, the density of fibrocytes, the density of interacting cells, the mean minimal distance between fibrocytes and $CD8^+$ T cells and the density of fibrocytes-$CD8^+$ T cells clusters.** Relationships between the density of $CD45^+$ $FSP1^+$ cells (**A**), the density of interacting cells (**B**), the mean minimal distance between fibrocytes and $CD8^+$ T cells (**C**), the density of mixed cell clusters (**D**) and the $FEV_1$/FVC ratio measured in control subjects (open circles) and COPD patients (black circles). The correlation coefficient (R) and significance level (P value) were obtained by using nonparametric Spearman analysis. n=20 specimens from control subjects, n=12 specimens from patients with COPD.

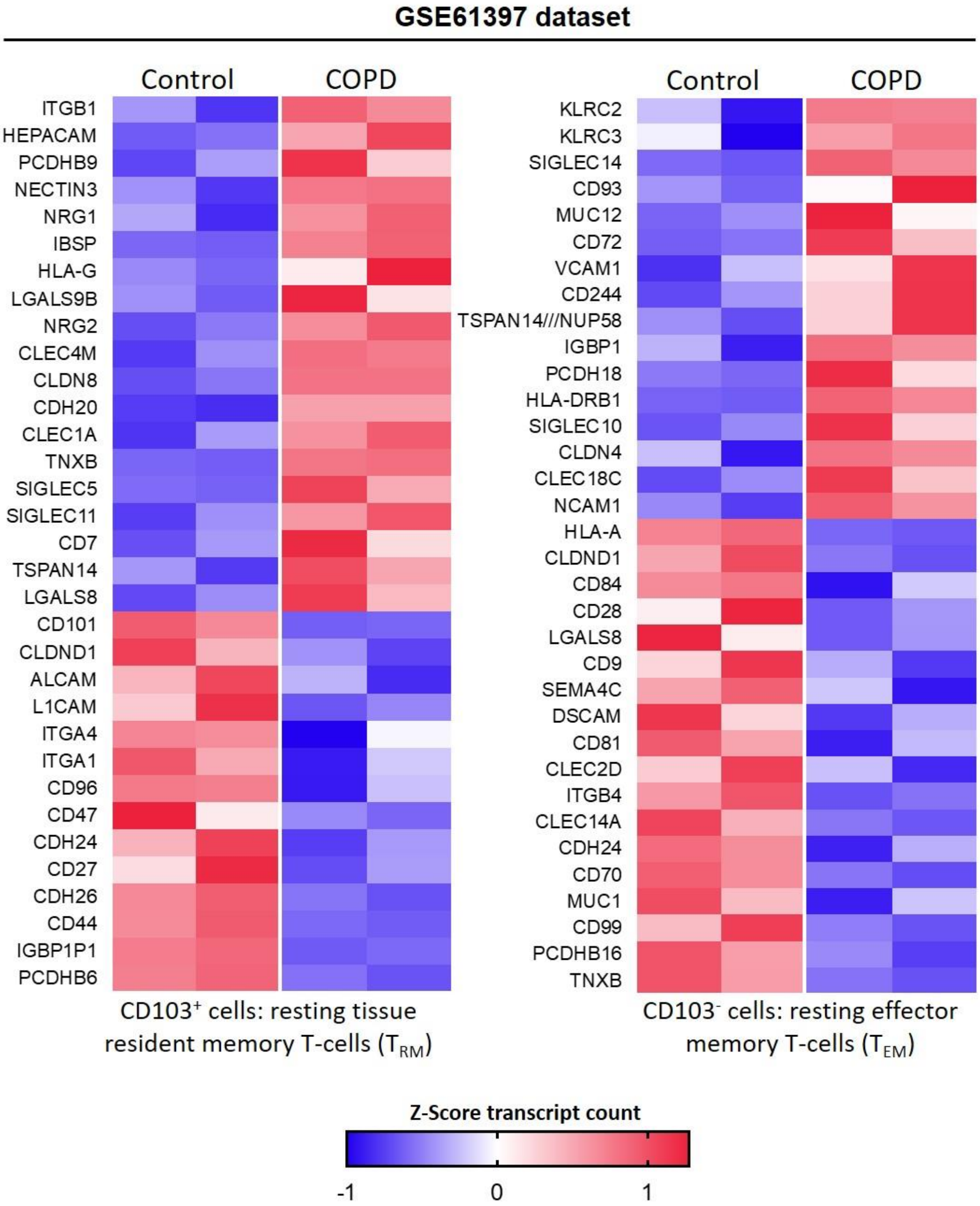


**Figure S5. Transcript levels of adhesion genes in CD8$^{+}$ T cells from COPD tissues compared to control tissues.** Heatmaps showing the expression of differentially expressed genes with p-value<0.05 of adhesion molecules and adhesion receptors in resting tissular tissue resident memory T-cells ($T_{RM}$) and effector memory T-cells ($T_{EM}$) from patients with COPD (n=2 independent samples) in comparison with control subjects (n=2 independent samples) (GEO accession GSE61397). Expression values are expressed as Z-score transformed transcript count.

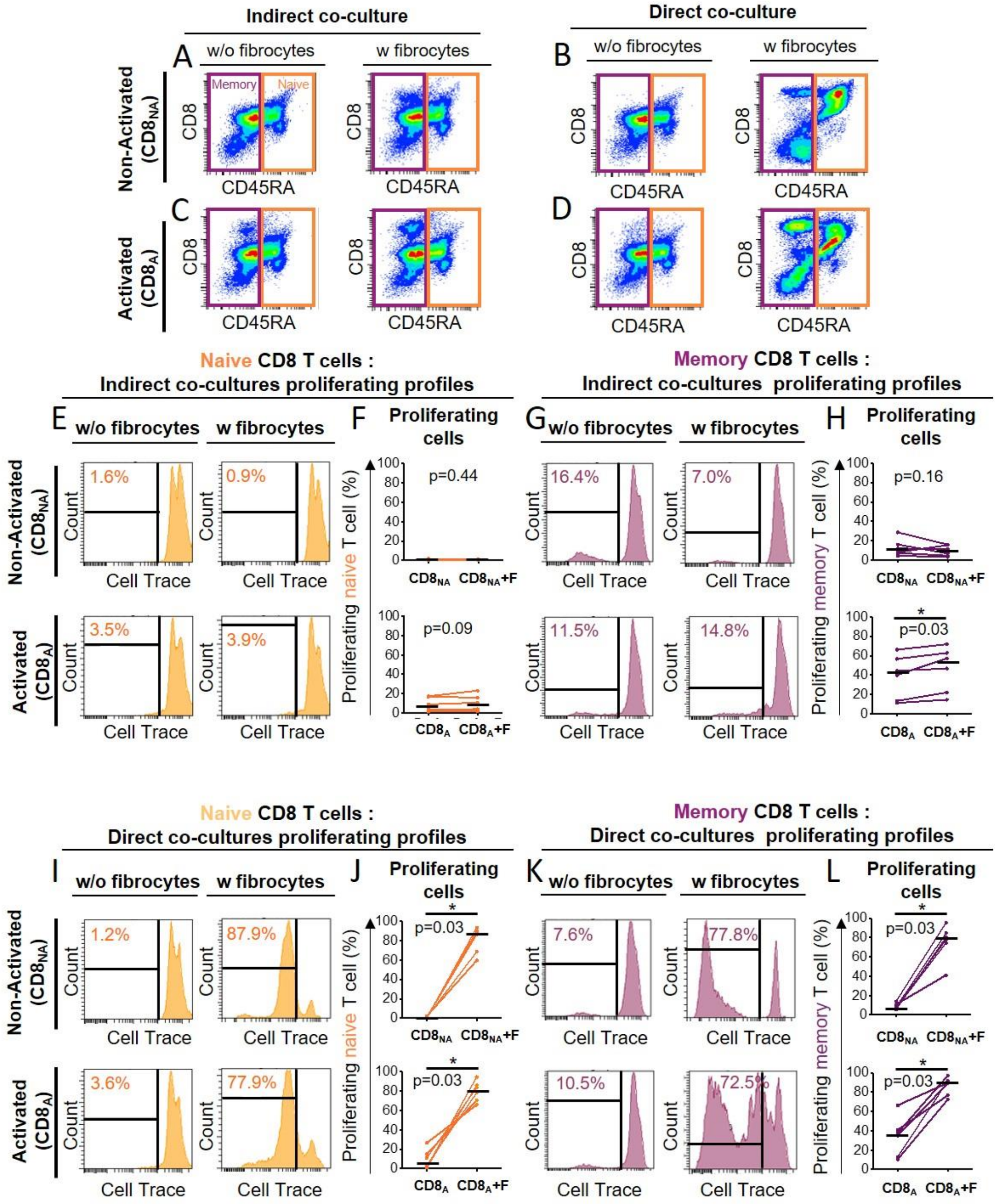


**Figure S6. Direct contact between fibrocytes and CD8$^+$ T cells preferentially increases proliferation of naïve CD8$^+$ T cells subset.** Prior to co-culture, CD8$^+$ T cells have been either non-activated ("CD8$_{NA}$") or activated ("CD8$_A$"). **(A**, **B**, **C**, **D)** Representative gating strategy for identification of naïve CD8$^+$CD45RA$^+$ cells and memory CD8$^+$CD45RA$^-$ CD8$^+$ T cells without (w/o) fibrocytes or with (w) fibrocytes in indirect (**A**, **C**) or direct (**B**, **D**) co-culture. Dot plots represent representative CD8-PerCP-Cy5-5 fluorescence (y-axis) versus CD45RA-FITC fluorescence (x-axis) of non-adherent cells removed from the culture. (**E**, **G**, **I**, **K**) Histograms represent representative cell count (y-axis) versus Cell Trace-Pacific Blue fluorescence (x-axis) in naïve CD8$^+$CD45RA$^+$ cells (**E**, **I**) and memory CD8$^+$CD45RA$^-$ cells (**G**, **K**) in co-culture without (w/o) fibrocytes or with (w) fibrocytes in indirect (**E**, **G**) or direct (**I**, **K**) co-culture. The distinct fluorescence peaks correspond to the different generations of CD8$^+$ T cells. The gate and the percentage indicate cells that have proliferated. (**F**, **H**, **J**, **L**) Comparison of quantifications of naïve (**F**, **J**) and memory (**H**, **L**) $CD8^+$ T cells that have proliferated, removed co-culture without fibrocyte (“CD8”) and with fibrocyte (“CD8+F”). n=6 independent experiments. Medians are represented as horizontal lines. * $P < 0.05$, Wilcoxon matched pairs test.

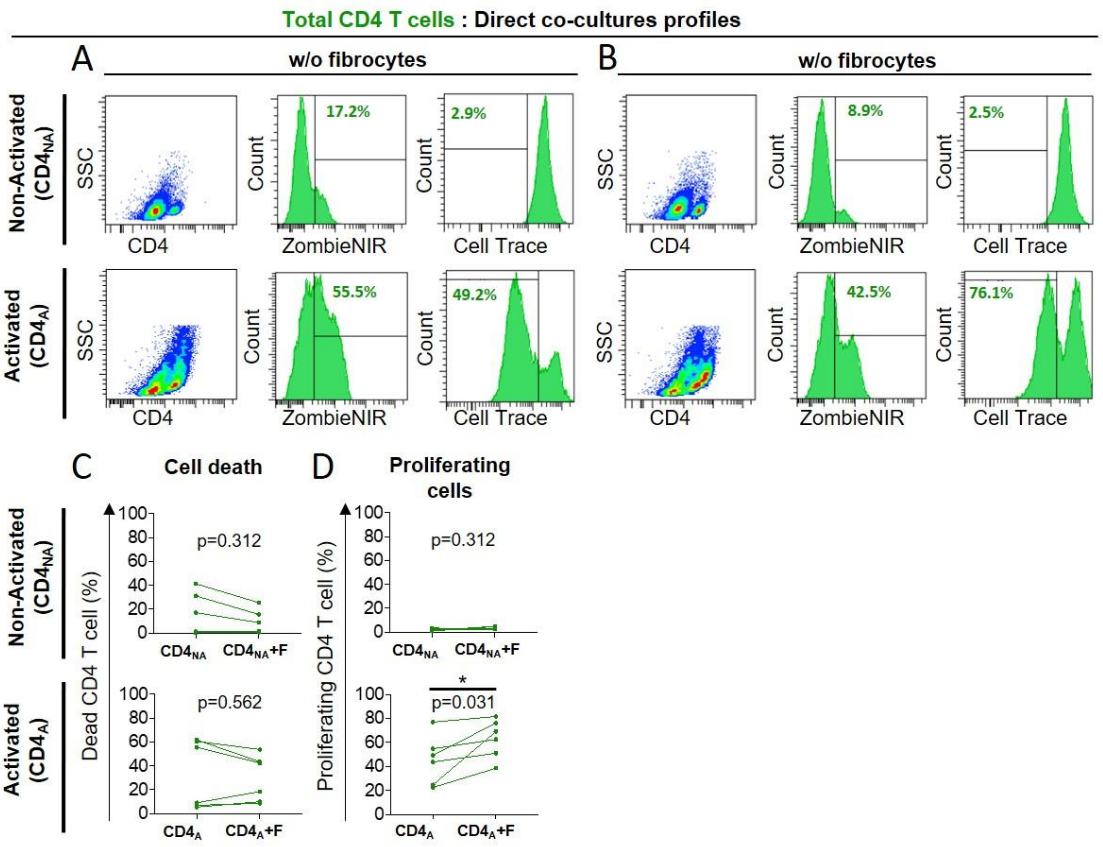


**Figure S7. CD4$^{+}$ T cells death and proliferation after 6 days in direct co-culture with fibrocytes.** Prior to co-culture, CD4$^{+}$ T cells have been either non-activated (“CD4$_{NA}$”) or activated (“CD4$_{A}$”). (**A, B**) Representative gating strategy for identification of CD4$^{+}$ T cells without (w/o) fibrocytes (**A**) or with (w) fibrocytes (**B**) in direct co-culture. Left panels: dot plots represent representative CD4-PerCP-Cy5-5 fluorescence (y-axis) versus side scatter (SSC, x-axis) of non-adherent cells removed from the culture. Mid panels: histograms represent representative cell count (y-axis) versus ZombieNIR-APC-Cy7 fluorescence (x-axis). Right panels: histograms represent representative cell count (y-axis) versus Cell Trace-Pacific Blue fluorescence (x-axis). The distinct fluorescence peaks correspond to the different generations of CD4$^{+}$ T cells. The gate and the percentage indicate cells that have proliferated. (**C**) Comparison of quantifications of dead CD4$^{+}$ T cells removed from co-culture without fibrocyte (“CD4”) and with fibrocyte (“CD4+F). (**D**) Comparison of quantifications of CD4$^{+}$ T cells that have proliferated, removed from co-culture. * $P < 0.05$, Wilcoxon matched paired tests.

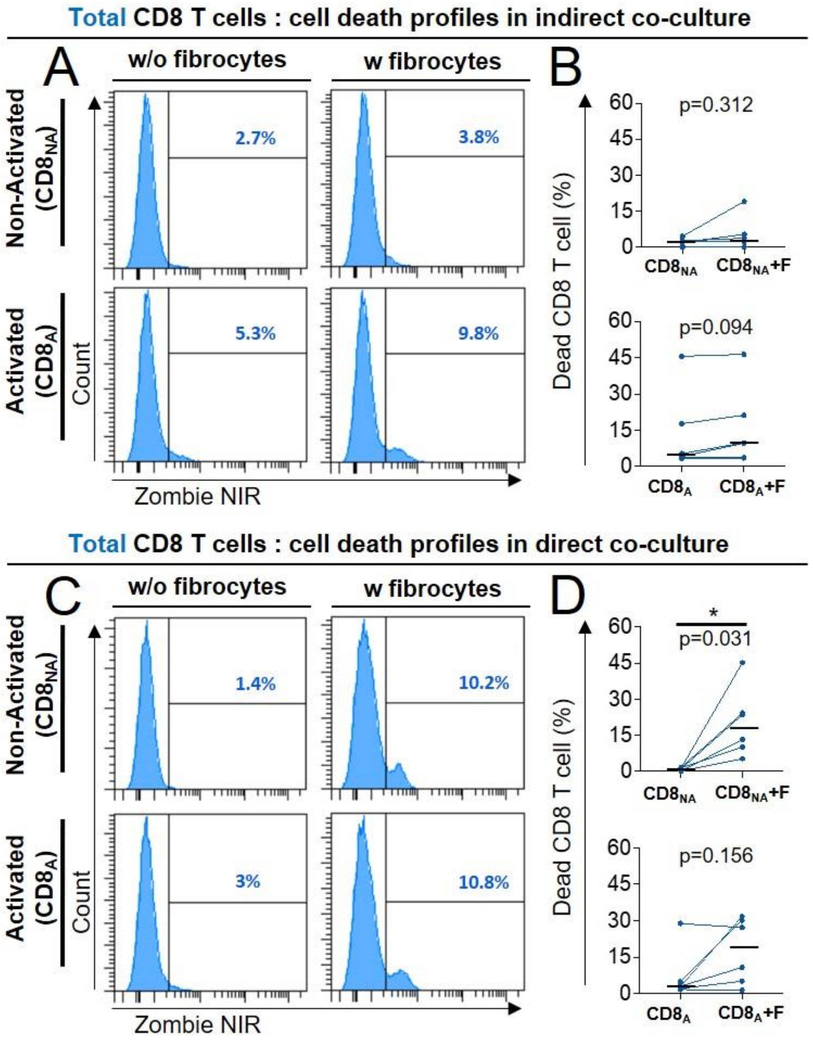


**Figure S8. CD8+ T cells death after 6 days of co-culture with fibrocytes.** Prior to co-culture, CD8+ T cells have been either non-activated ("$CD8_{NA}$") or activated ("$CD8_A$"). (**A**, **C**) Representative gating strategy for identification of dead CD8+ T cells without (w/o) fibrocytes (left panels) or with (w) fibrocytes (right panels) in indirect (**A**) or direct (**C**) co-culture. Histograms represent representative cell count (y-axis) versus Zombie NIR fluorescence (x-axis). (**B**, **D**) Comparison of quantifications of dead CD8+ T cells removed from co-culture without fibrocyte ("CD8") and with fibrocyte ("CD8+F") in indirect (**B**) or direct (**D**) co-culture. Medians are represented as horizontal lines. * $P < 0.05$, Wilcoxon matched paired tests.

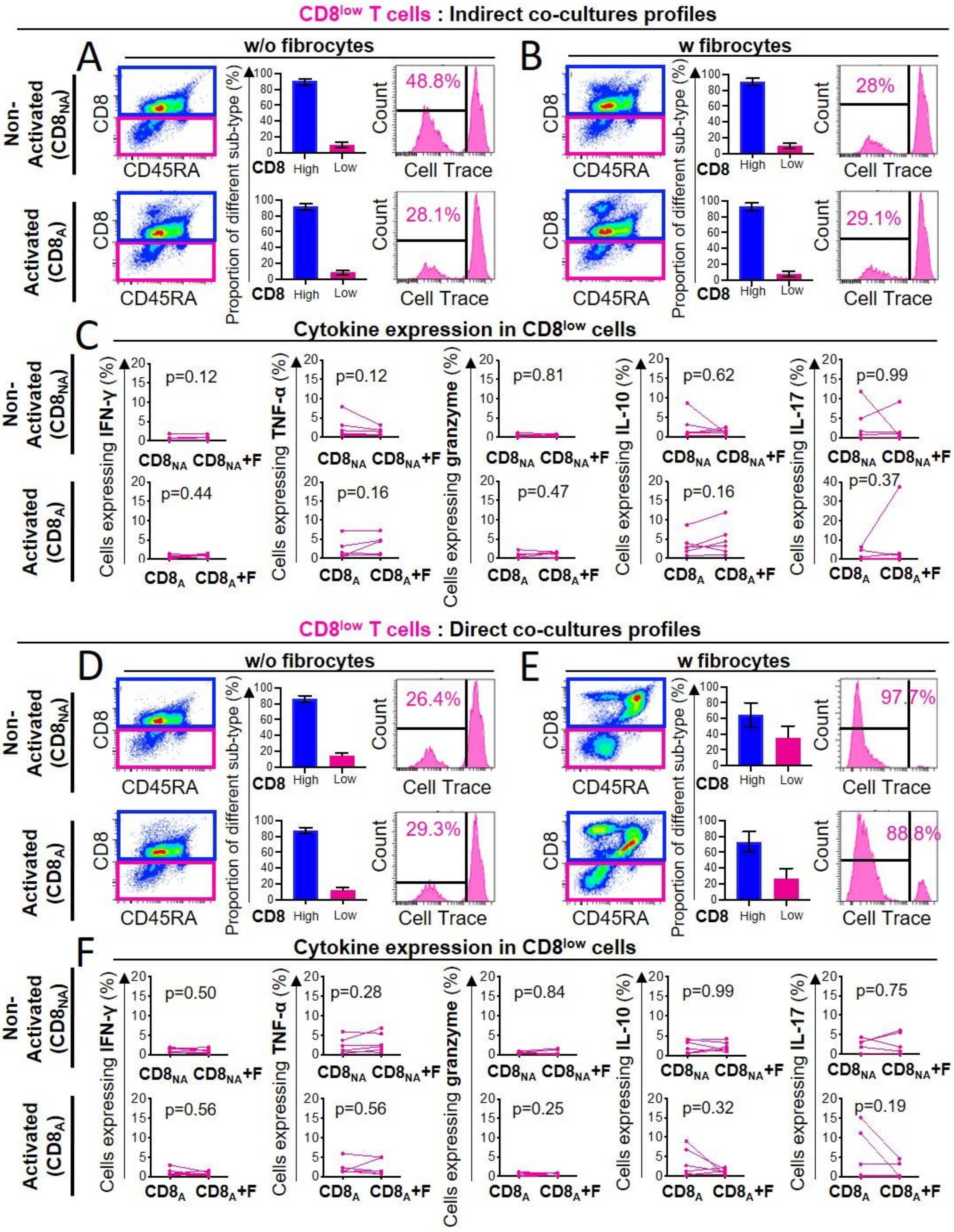


**Figure S9. Direct contact between fibrocytes and CD8$^+$ T cells promote phenotypic differences in CD8 expression.** Prior to co-culture, CD8$^+$ T cells have been either non-activated ("CD8$_{NA}$") or activated ("CD8$_A$"). (**A**, **B**, **D**, **E**) Representative gating strategy for identification of differents $CD8^+$ T cells sub-types without (w/o) fibrocytes (**A**, **D**) or with (w) fibrocytes (**B**, **E**) in indirect (**A**, **B**) or direct (**D**, **E**) co-culture. Left panels: dot plots represent representative CD8-PerCP-Cy5-5 fluorescence (y-axis) versus CD45RA-FITC fluorescence (x-axis) of non-adherent cells removed from the culture. $CD8^+$ T cells that express respectively low and high levels of CD8 are gated respectively in pink ($CD8^{low}$) and blue ($CD8^{high}$). Middle panels: histograms represent the proportion of the two differents sub-types ($CD8^{low}$ and $CD8^{high}$) in the total population. Right histograms represent representative cell count (y-axis) versus Cell Trace-Pacific Blue fluorescence (x-axis) for the $CD8^{low}$ population sub-type. The distinct fluorescence peaks correspond to the different generations of $CD8^+$ T cells. The gate and the percentage indicate cells that have proliferated. (**C**, **F**) Quantifications of $CD8^{low}$ T cells expressing IFN-γ, TNF-α, Granzyme, IL-10, IL-17 for indirect co-culture (**C**) and direct co-culture (**F**) and without ($CD8_{NA}$ or $CD8_{A}$) or with ($CD8_{NA}$+F or CD8A+F ) fibrocytes. n=6 independent experiments. * $P < 0.05$, Wilcoxon matched pairs test.

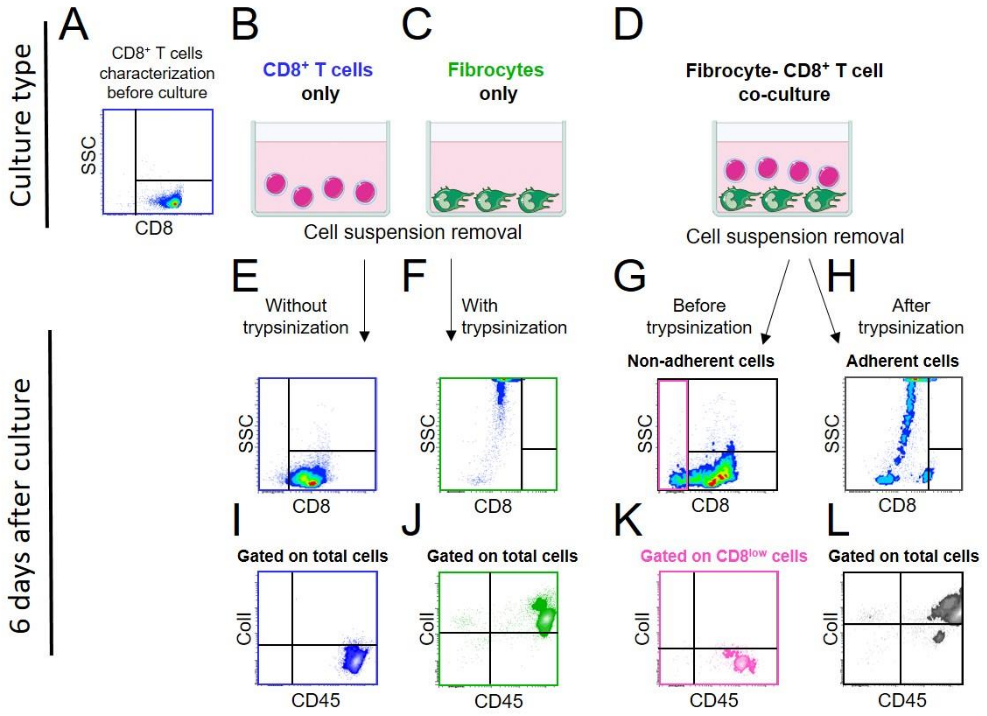


**Figure S10: The CD8$^{low}$ population appears in co-culture and is distinct from the CD45$^{+}$ Collagen I$^{+}$ population.** Prior to co-culture, CD8$^{+}$ T cells have been activated. (**A**, **B**, **C**, **D**) Experiment design. Pure CD8$^{+}$ T cells were characterized by flow cytometry for CD8 expression (**A**) before being were cultured either alone (**B**) or with fibrocytes (**D**). Fibrocytes were either cultured alone (**C**) or with CD8$^{+}$ T cells (**D**). (**E**, **F**, **G**, **H**) Dot plots represent representative CD8-PerCP-Cy5-5 fluorescence (y-axis) versus granularity (Side Scatter, SSC) (x-axis) of cells removed either without trypsinization (**E**, **G**) or with trypsinization (**F**, **H**) after 6 days in culture/co-culture. (**I**, **J**, **K**, **L**) Dot plots represent representative Collagen Type I (ColI)-FITC fluorescence (y-axis) versus CD45-APC fluorescence (x-axis) of cells removed either without trypsinization (**I**, **K**) or with trypsinization (**J**, **L**) after 6 days in culture/co-culture. For conditions with CD8$^{+}$ T cells only (**I**), fibrocytes only (**J**), and adherent cells removed from co-culture (**K**), the cytographs were generated by gating the total cell population. For non-adherent cells removed from co-culture (**L**), the cytograph was generated by gating the CD8$^{low}$ population (pink).

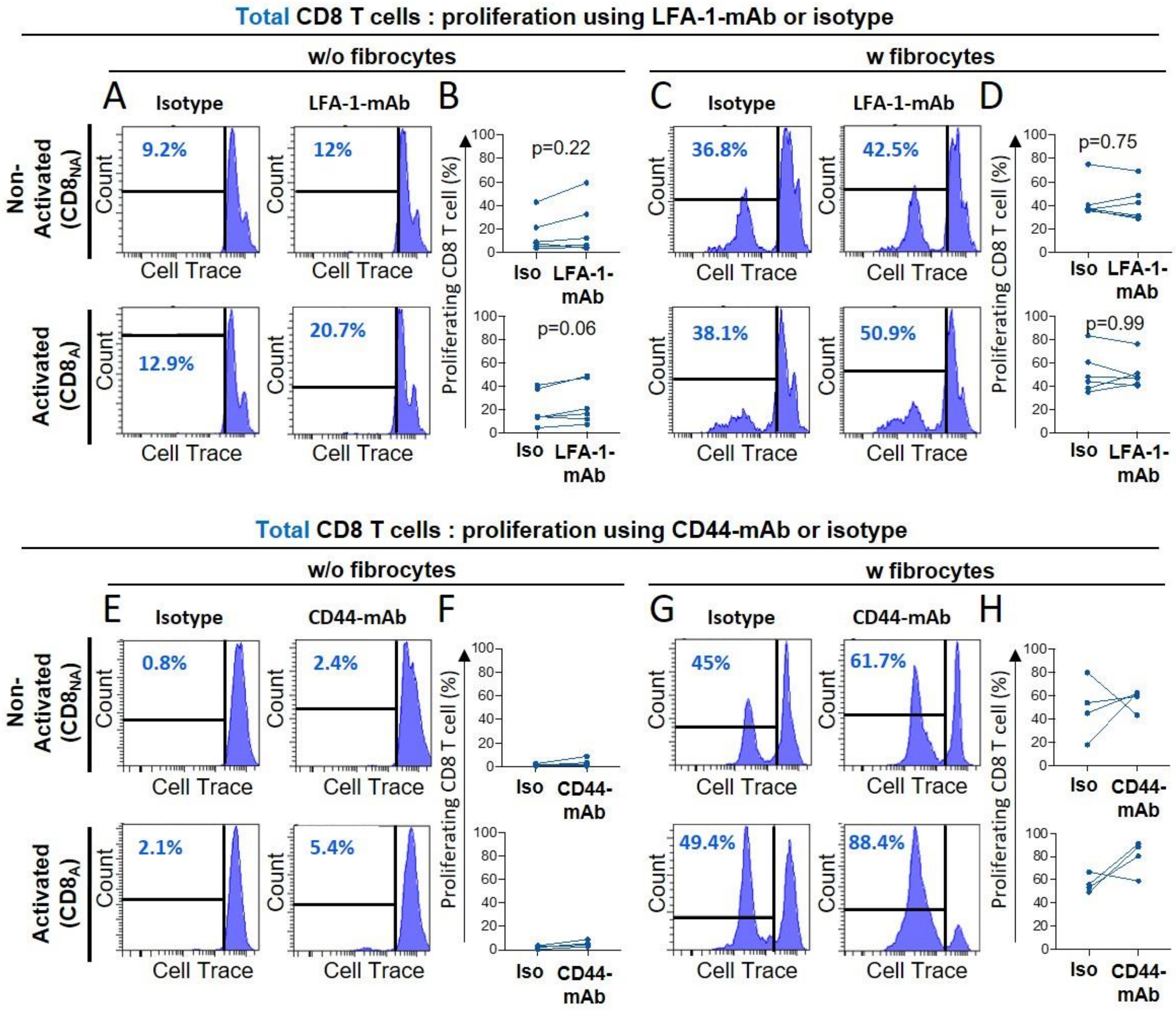


**Figure S11: LFA-1 or CD44 blockade is not sufficient to decrease $CD8^+$ T cells proliferation induction.** Prior to co-culture, $CD8^+$ T cells have been either non-activated (“$CD8_{NA}$”) or activated (“$CD8_A$”). **(A**, **C**, **E**, **G**) Representative gating strategy for identification of proliferating $CD8^+$ T cells without (w/o) fibrocytes (**A**, **E**) or with (w) fibrocytes (**C**, **G**) using neutralizing LFA-1-mAb (**A**, **C**) or neutralizing CD44-mAb (**E**, **G**) and respective control isotype. Histograms represent representative cell count (y-axis) versus Cell Trace-Pacific Blue fluorescence (x-axis). The distinct fluorescence peaks correspond to the different generations of $CD8^+$ T cells. The gate and the percentage indicate cells that have proliferated. (**B**, **D**, **F**, **H**) Comparison of quantifications of $CD8^+$ T cells that have proliferated, removed from co-culture treated with neutralizing LFA-1-mAb (**B**, **D**) or neutralizing CD44-mAb (**F**, **H**) and respective control isotype. (**A-D**) n=6 independent experiments. * $P < 0.05$, Wilcoxon matched pairs test. (**E-H**) n=4 independent experiments.

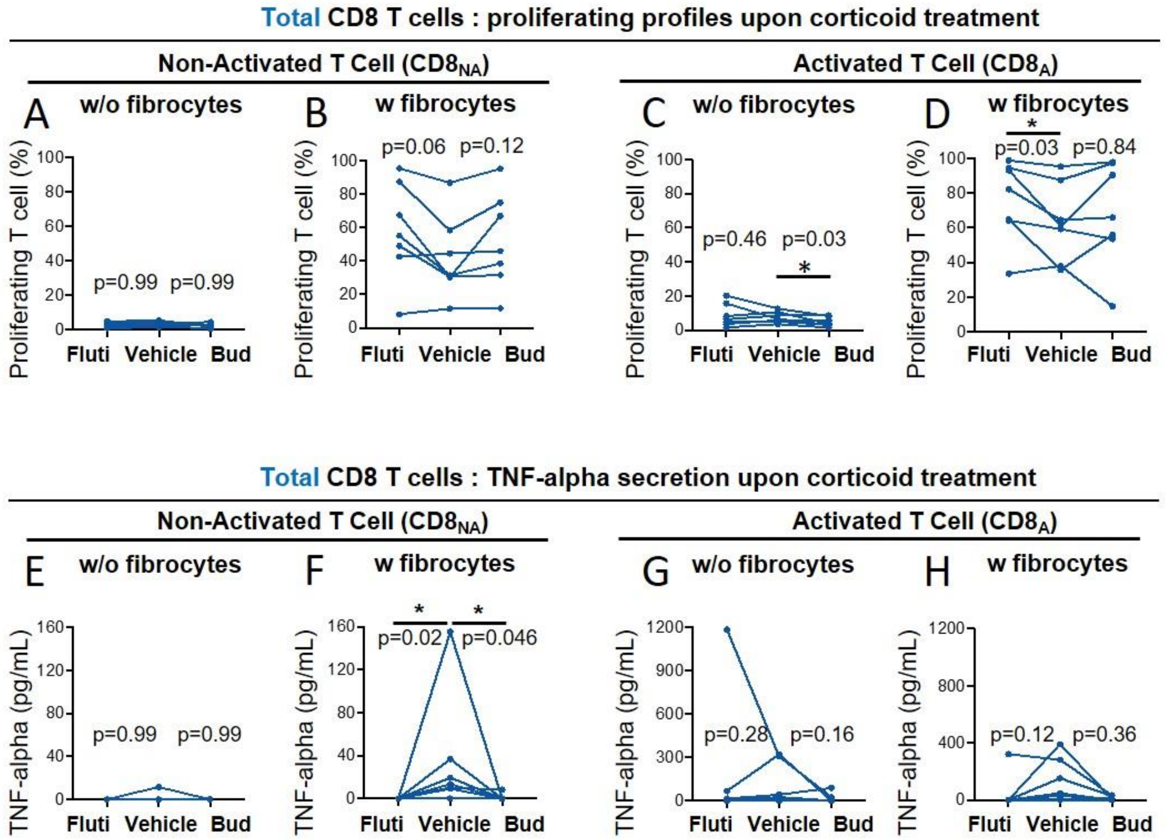


**Figure S12: Glucocorticoid drugs significantly decrease fibrocyte-induced TNF-α secretion by CD8⁺ T cells but not the proliferation induction.** Prior to co-culture, $CD8^+$ T cells have been either non-activated or activated. (**A**, **B**, **C**, **D**) Comparison of quantifications of $CD8^+$ T cells that have proliferated, removed from co-culture without fibrocytes (w/o fibrocytes) (**A**, **C**) and with fibrocytes (w fibrocytes) (**B**, **D**) for non-activated $CD8^+$ T cells (**A**, **B**) or activated ones (**C**, **D**) and using fluticasone propionate (Fluti), budesonide (Bud) or vehicle. (**E**, **F**, **G**, **H**) Secreted TNF-α concentrations in supernatants from co-cultures without fibrocytes (w/o fibrocytes) (**E**, **G**) and with fibrocyte (w fibrocytes) (**F**, **H**) for non-activated $CD8^+$ T cells (**E**, **F**) or activated ones (**G**, **H**) and using fluticasone propionate (Fluti), budesonide (Bud) or vehicle. Values below the detection limit were counted as zero. n=6 independent experiments. * $P < 0.05$, Friedman test.

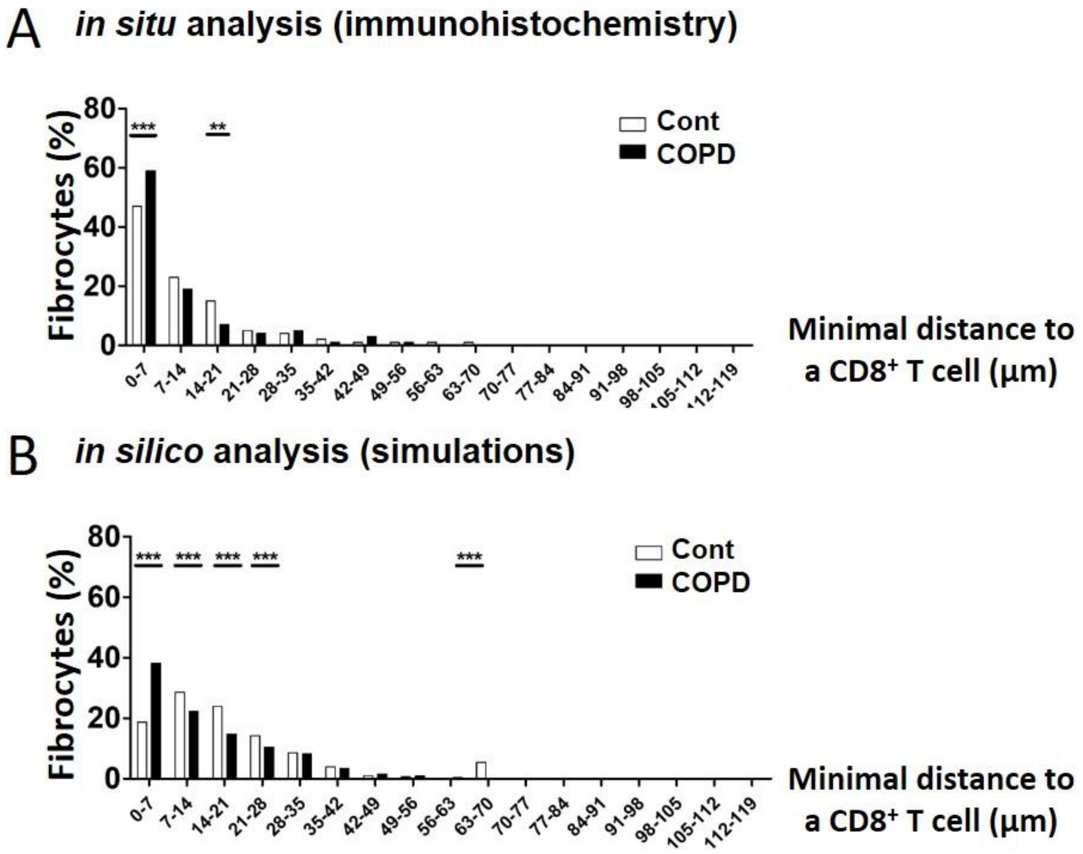


**Figure S13: Minimal intercellular distance distributions are similar between *in situ* analyses and simulations.** Mean frequency distributions of minimal distances (with 7 µm binning) between fibrocytes and CD8$^+$ T cells for control subjects (white) and patients with COPD (black), built from in situ analyses (**A**) and final state of simulations (**B**). Error bars indicate standard error of the mean. Two-way ANOVA for repeated measures with Bonferroni post-tests. $F(16, 493)=3.2$, $P<0.0001$ (**A**); $F(16, 5406)=140$, $P<0.0001$ (**B**). P-values of with Bonferroni post-tests are indicated on the figure with the following symbols: **: $P<0.01$, ***: $P<0.001$.

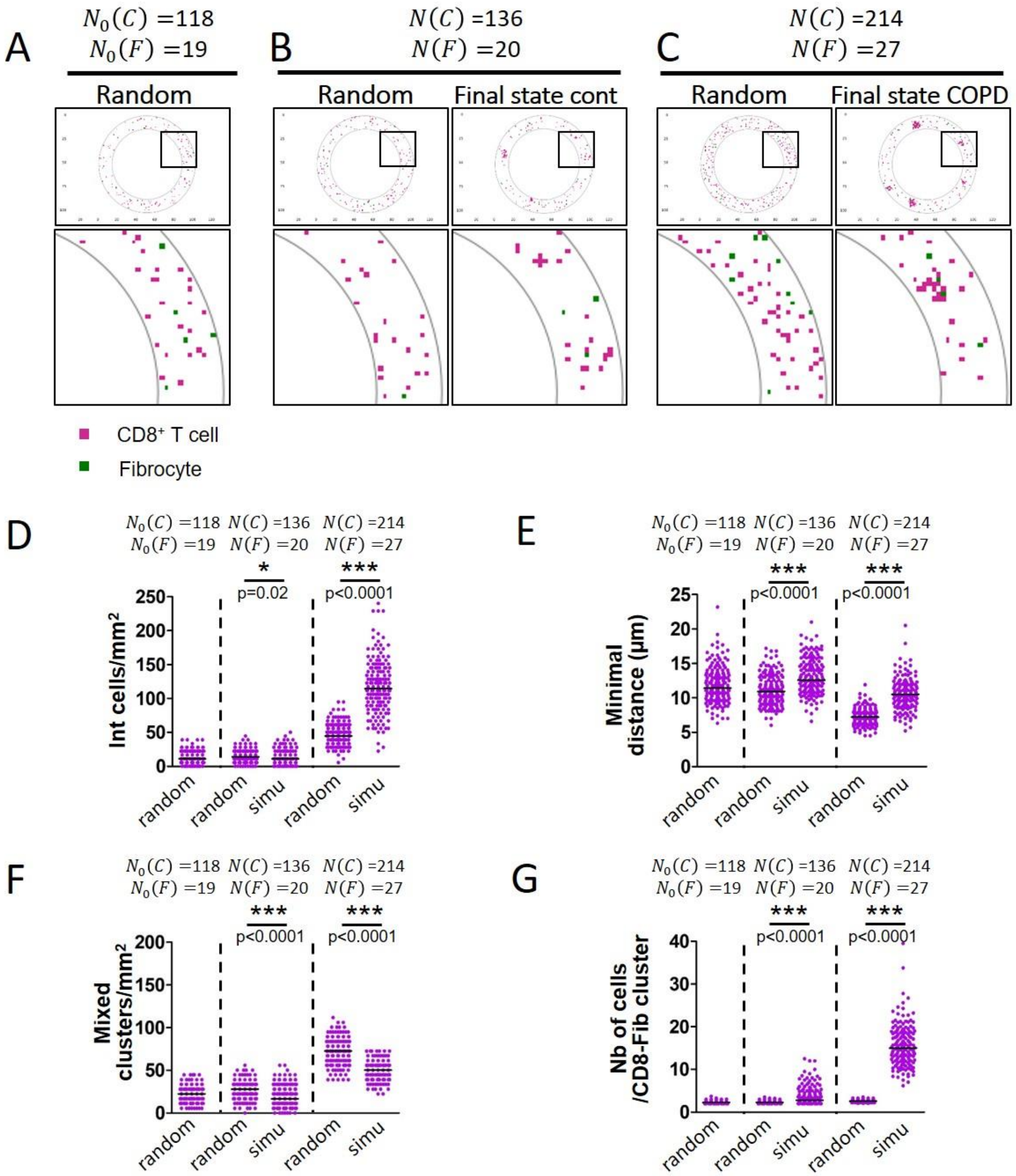


**Figure S14: Spatial cellular repartition of cells obtained at the final state of simulations are distinct from random distributions.** Final states of the simulations obtained after 20 years of control dynamics and COPD dynamics are compared to random distributions with indicated fibrocyte and CD8+ T cells numbers. (**A**) Selected representative picture of the peribronchial area with $N_0(C)$ =118 CD8+ T cells and $N_0(F)$ =19 fibrocytes randomly seeded (initial state for simulations). (**B**) Selected representative pictures of the peribronchial area with $N(C)$ =136 CD8+ T cells and $N(F)$ =20 fibrocytes. Left panel: random distribution. Right panel: example of distribution obtained after 20 years of control dynamics. (**C**) Selected representative pictures of the peribronchial area with $N(C)$ =214 CD8$^+$ T cells and $N(F)$ =37 fibrocytes. Left panel: random distribution. Right panel: example of distribution obtained after 20 years of COPD dynamics. (**A-C**), Images surrounded by black squares: higher magnifications of peribronchial area. CD8$^+$ T cells and fibrocytes are represented respectively by pink and green squares. (**D**), CD8$^+$ T cells (left) and fibrocyte (right) densities. (**E**) Interacting cells densities of interacting cells. (**F**) Mean minimal distances between fibrocyte and CD8$^+$ T cells. (**G**) CD8$^+$ T cells-fibrocytes-containing clusters ("CD8-Fib clusters") densities. (**H**) mean number of cells by CD8-Fib clusters. (**C**-**G**) n=160 simulations for each situation. The medians are represented as horizontal lines. *: P<0.05, ***: P<0.001. unpaired t-tests or Mann-Whitney tests.

**Table S1. Association between density of fibrocytes and clinical characteristics**

| | Density of fibrocytes | | |
|---|---|---|---|
| | **Spearman r** | **95% confidence interval** | **P value** |
| Age (yrs.) | -0,14 | [ -0.48 to 0.23] | 0,43 |
| Body-mass index (kg/$m^2$) | -0,12 | [ -0.46 to 0.25] | 0,51 |
| Pack years (no.) | 0,14 | [ -0.24 to 0.48] | 0,46 |
| **LFT** | | | |
| $FEV_1$ (% pred.) | -0,44 | [-0.69 to -0.10] | **0,01** |
| $FEV_1$/FVC ratio (%) | -0,38 | [-0.65 to -0.03] | **0,03** |
| FVC (% pred.) | -0,38 | [-0.65 to -0.03] | **0,03** |
| RV (% pred.) | 0,30 | [-0.07 to 0.59] | 0,10 |
| TLCO (% pred.) | -0,56 | [-0.78 to -0.24] | **0,001** |
| **Six-minute walk test distance (m)** | -0,14 | [-0.52 to 0.29] | 0,52 |
| **Arterial blood gases** | | | |
| $PaO_2$ (mm Hg) | 0,08 | [-0.30 to 0.44] | 0,67 |
| $PaCO_2$ (mm Hg) | 0,26 | [-0.13 to 0.57] | 0,17 |
| **CT parameters** | | | |
| Bronchi: | | | |
| WA4% | 0,08 | [-0.33 to 0.46] | 0,71 |
| WT4 (mm) | 0,03 | [-0.37 to 0.42] | 0,88 |
| WA5% | 0,10 | [-0.31 to 0.48] | 0,62 |
| WT5 (mm) | 0,19 | [-0.23 to 0.55] | 0,36 |
| Emphysema: | | | |
| LAA (%) | 0,40 | [0.04 to 0.67] | **0,03** |
| Air trapping: | | | |
| MLA E (HU) | -0,38 | [-0.69 to 0.04] | 0,07 |
| MLA I (HU) | -0,36 | [-0.65 to 0.02] | 0,053 |
| MLA I-E (HU) | -0,15 | [-0.53 to 0.28] | 0,48 |
| Pulmonary Vessels | | | |
| $\%CSA_{<5}$ | -0,41 | [-0.68 to -0.05] | **0,02** |
| $\%CSA_{5-10}$ | -0,22 | [-0.55 to 0.16] | 0,24 |
| $CSN_{<5}$ | -0,38 | [-0.66 to -0.02] | **0,04** |
| $CSN_{5-10}$ | -0,41 | [-0.68 to -0.05] | **0,02** |

$FEV_1$, forced expiratory volume in 1 second; FVC, forced vital capacity; LFT, lung function test; RV, residual volume; TLCO, Transfer Lung capacity of Carbon monoxide, $PaO_2$, partial arterial oxygen pressure, $PaCO_2$, partial arterial carbon dioxide pressure; WA, mean wall area; LA, mean lumen area, WA%, mean wall area percentage; WT, wall thickness; LAA, low-attenuation area; MLA E or I, mean lung attenuation value during expiration or inspiration. MLA I-E, difference between inspiratory and expiratory mean lung attenuation value. $\%CSA_{<5}$, percentage of total lung area taken up by the cross-sectional area of pulmonary vessels less than 5 $mm^2$; $\%CSA_{5-10}$, percentage of total lung area taken up by the cross-sectional area of pulmonary vessels between 5 and 10 $mm^2$; $CSN_{<5}$, number of vessels less than 5 $mm^2$ normalized by total lung area; $CSN_{5-10}$, number of vessels between 5 and 10 $mm^2$ normalized by total lung area; NR: not relevant. The correlation coefficient (r), 95% confidence interval and significance level (P value), were obtained by using nonparametric Spearman analysis.

**Table S2. Association between density of CD8+ T cells and clinical characteristics**

| | Density of CD8+ T cells | | |
|---|---|---|---|
| | **Spearman r** | **95% confidence interval** | **P value** |
| Age (yrs.) | -0,09 | [-0.43 to 0.28] | 0,63 |
| Body-mass index ($kg/m^2$) | -0,07 | [-0.42 to 0.30] | 0,71 |
| Pack years (no.) | 0,33 | [-0.04 to 0.62] | 0,07 |
| **LFT** | | | |
| $FEV_1$ (% pred.) | -0,25 | [ -0.56 to 0.12] | 0,16 |
| $FEV_1$/FVC ratio (%) | -0,30 | [ -0.60 to 0.06] | 0,09 |
| FVC (% pred.) | -0,18 | [ -0.50 to 0.19] | 0,33 |
| RV (% pred.) | 0,24 | [ -0.13 to 0.55] | 0,19 |
| TLCO (% pred.) | -0,25 | [ -0.57 to 0.14] | 0,19 |
| **Six-minute walk test distance (m)** | -0,06 | [ -0.46 to 0.37] | 0,79 |
| **Arterial blood gases** | | | |
| $PaO_2$ (mm Hg) | 0,02 | [-0.35 to 0.39] | 0,91 |
| $PaCO_2$ (mm Hg) | 0,40 | [ 0.04 to 0.67] | **0,03** |
| **CT parameters** | | | |
| Bronchi: | | | |
| WA4% | 0,35 | [ -0.05 to 0.66] | 0,08 |
| WT4 (mm) | 0,31 | [ -0.09 to 0.63] | 0,12 |
| WA5% | 0,32 | [ -0.09 to 0.64] | 0,11 |
| WT5 (mm) | 0,31 | [ -0.10 to 0.63] | 0,12 |
| Emphysema: | | | |
| LAA (%) | 0,39 | [ 0.02 to 0.66] | **0,03** |
| Air trapping: | | | |
| MLA E (HU) | -0,49 | [ -0.75 to -0.09] | **0,02** |
| MLA I (HU) | -0,48 | [ -0.73 to -0.13] | **0,008** |
| MLA I-E (HU) | 0,15 | [ -0.28 to 0.53] | 0,48 |
| Pulmonary Vessels | | | |
| $\%CSA_{<5}$ | -0,38 | [ -0.66 to -0.01] | **0,04** |
| $\%CSA_{5-10}$ | -0,44 | [ -0.70 to -0.08] | **0,02** |
| $CSN_{<5}$ | -0,29 | [ -0.59 to 0.09] | 0,12 |
| $CSN_{5-10}$ | -0,41 | [ -0.68 to -0.05] | **0,02** |

$FEV_1$, forced expiratory volume in 1 second; FVC, forced vital capacity; LFT, lung function test; RV, residual volume; TLCO, Transfer Lung capacity of Carbon monoxide, $PaO_2$, partial arterial oxygen pressure, $PaCO_2$, partial arterial carbon dioxide pressure; WA, mean wall area; LA, mean lumen area, WA%, mean wall area percentage; WT, wall thickness; LAA, low-attenuation area; MLA E or I, mean lung attenuation value during expiration or inspiration. MLA I-E, difference between inspiratory and expiratory mean lung attenuation value. $\%CSA_{<5}$, percentage of total lung area taken up by the cross-sectional area of pulmonary vessels less than 5 $mm^2$; $\%CSA_{5–10}$, percentage of total lung area taken up by the cross-sectional area of pulmonary vessels between 5 and 10 $mm^2$; $CSN_{<5}$, number of vessels less than 5 $mm^2$ normalized by total lung area; $CSN_{5-10}$, number of vessels between 5 and 10 $mm^2$ normalized by total lung area; NR: not relevant. The correlation coefficient (r), 95% confidence interval and significance level (P value), were obtained by using nonparametric Spearman analysis.

**Table S3. Association between interacting cell density and clinical characteristics**

| | Interacting cells density (D8) | | |
|---|---|---|---|
| | **Spearman r** | **95% confidence interval** | **P value** |
| Age (yrs.) | -0,17 | [-0.50 to 0.20] | 0,34 |
| Body-mass index (kg/m$^2$) | -0,13 | [-0.47 to 0.24] | 0,48 |
| Pack years (no.) | 0,22 | [-0.16 to 0.55] | 0,23 |
| **LFT** | | | |
| $FEV_1$ (% pred.) | -0,38 | [-0.65 to -0.02] | **0,03** |
| $FEV_1$/FVC ratio (%) | -0,37 | [-0.64 to -0.02] | **0,04** |
| FVC (% pred.) | -0,27 | [-0.57 to 0.10] | 0,13 |
| RV (% pred.) | 0,15 | [-0.22 to 0.48] | 0,41 |
| TLCO (% pred.) | -0,50 | [-0.74 to -0.16] | **0,005** |
| **Six-minute walk test distance (m)** | -0,25 | [-0.60 to 0.18] | 0,24 |
| **Arterial blood gases** | | | |
| $PaO_2$ (mm Hg) | 0,25 | [-0.13 to 0.57] | 0,18 |
| $PaCO_2$ (mm Hg) | 0,36 | [-0.01 to 0.64] | 0,05 |
| **CT parameters** | | | |
| Bronchi: | | | |
| WA4% | 0,04 | [-0.36 to 0.43] | 0,83 |
| WT4 (mm) | 0,02 | [-0.38 to 0.41] | 0,93 |
| WA5% | 0,11 | [-0.30 to 0.49] | 0,58 |
| WT5 (mm) | 0,17 | [-0.24 to 0.53] | 0,41 |
| Emphysema: | | | |
| LAA (%) | 0,30 | [-0.08 to 0.60] | 0,11 |
| Air trapping: | | | |
| MLA E (HU) | -0,39 | [-0.69 to 0.026] | 0,06 |
| MLA I (HU) | -0,38 | [-0.66 to 0.001] | **0,04** |
| MLA I-E (HU) | 0,02 | [-0.40 to 0.43] | 0,92 |
| Pulmonary Vessels | | | |
| $\%CSA_{<5}$ | -0,39 | [-0.66 to -0.02] | **0,03** |
| $\%CSA_{5-10}$ | -0,22 | [-0.54 to 0.16] | 0,24 |
| $CSN_{<5}$ | -0,32 | [-0.61 to 0.06] | 0,09 |
| $CSN_{5-10}$ | -0,39 | [-0.66 to -0.02] | **0,03** |

$FEV_1$, forced expiratory volume in 1 second; FVC, forced vital capacity; LFT, lung function test; RV, residual volume; TLCO, Transfer Lung capacity of Carbon monoxide, $PaO_2$, partial arterial oxygen pressure, $PaCO_2$, partial arterial carbon dioxide pressure; WA, mean wall area; LA, mean lumen area, WA%, mean wall area percentage; WT, wall thickness; LAA, low-attenuation area; MLA E or I, mean lung attenuation value during expiration or inspiration. MLA I-E, difference between inspiratory and expiratory mean lung attenuation value. $\%CSA_{<5}$, percentage of total lung area taken up by the cross-sectional area of pulmonary vessels less than 5 $mm^2$; $\%CSA_{5–10}$, percentage of total lung area taken up by the cross-sectional area of pulmonary vessels between 5 and 10 $mm^2$; $CSN_{<5}$, number of vessels less than 5 $mm^2$ normalized by total lung area; $CSN_{5-10}$, number of vessels between 5 and 10 $mm^2$ normalized by total lung area; NR: not relevant. The correlation coefficient (r), 95% confidence interval and significance level (P value), were obtained by using nonparametric Spearman analysis.

**Table S4. Association between the mean minimal distance between fibrocytes and CD8$^{+}$ T cells and clinical characteristics**

| | Mean minimal distance between fibrocytes and CD8$^{+}$ T cells | | |
|---|---|---|---|
| | **Spearman r** | **95% confidence interval** | **P value** |
| Age (yrs.) | -0,02 | [-0.38 to 0.35] | 0,92 |
| Body-mass index (kg/m$^2$) | -0,10 | [-0.45 to 0.27] | 0,58 |
| Pack years (no.) | -0,10 | [-0.46 to 0.29] | 0,62 |
| **LFT** | | | |
| $FEV_1$ (% pred.) | 0,24 | [-0.13 to 0.56] | 0,19 |
| $FEV_1$/FVC ratio (%) | 0,42 | [0.06 to 0.68] | **0,02** |
| FVC (% pred.) | 0,10 | [-0.27 to 0.45] | 0,59 |
| RV (% pred.) | -0,14 | [-0.48 to 0.24] | 0,45 |
| TLCO (% pred.) | 0,27 | [-0.12 to 0.59] | 0,16 |
| **Six-minute walk test distance (m)** | -0,02 | [-0.44 to 0.41] | 0,92 |
| **Arterial blood gases** | | | |
| $PaO_2$ (mm Hg) | -0,05 | [-0.42 to 0.33] | 0,79 |
| $PaCO_2$ (mm Hg) | -0,45 | [-0.70 to -0.09] | **0,01** |
| **CT parameters** | | | |
| Bronchi: | | | |
| WA4% | -0,31 | [-0.64 to 0.11] | 0,13 |
| WT4 (mm) | -0,24 | [-0.59 to 0.18] | 0,24 |
| WA5% | -0,52 | [-0.76 to -0.14] | **0,008** |
| WT5 (mm) | -0,50 | [-0.75 to -0.12] | **0,01** |
| Emphysema: | | | |
| LAA (%) | -0,36 | [-0.65 to 0.02] | 0,06 |
| Air trapping: | | | |
| MLA E (HU) | 0,52 | [0.12 to 0.77] | **0,01** |
| MLA I (HU) | 0,50 | [0.15 to 0.74] | **0,006** |
| MLA I-E (HU) | -0,19 | [-0.57 to 0.25] | 0,38 |
| Pulmonary Vessels | | | |
| $\%CSA_{<5}$ | 0,19 | [-0.20 to 0.53] | 0,32 |
| $\%CSA_{5-10}$ | 0,29 | [-0.10 to 0.60] | 0,13 |
| $CSN_{<5}$ | 0,15 | [-0.24 to 0.59] | 0,43 |
| $CSN_{5-10}$ | 0,26 | [-0.14 to 0.57) | 0,20 |

$FEV_1$, forced expiratory volume in 1 second; FVC, forced vital capacity; LFT, lung function test; RV, residual volume; TLCO, Transfer Lung capacity of Carbon monoxide, $PaO_2$, partial arterial oxygen pressure, $PaCO_2$, partial arterial carbon dioxide pressure; WA, mean wall area; LA, mean lumen area, WA%, mean wall area percentage; WT, wall thickness; LAA, low-attenuation area; MLA E or I, mean lung attenuation value during expiration or inspiration. MLA I-E, difference between inspiratory and expiratory mean lung attenuation value. $\%CSA_{<5}$, percentage of total lung area taken up by the cross-sectional area of pulmonary vessels less than 5 $mm^2$; $\%CSA_{5–10}$, percentage of total lung area taken up by the cross-sectional area of pulmonary vessels between 5 and 10 $mm^2$; $CSN_{<5}$, number of vessels less than 5 $mm^2$ normalized by total lung area; $CSN_{5-10}$, number of vessels between 5 and 10 $mm^2$ normalized by total lung area; NR: not relevant. The correlation coefficient (r), 95% confidence interval and significance level (P value), were obtained by using nonparametric Spearman analysis.

**Table S5. Association between the density of mixed cell clusters and clinical characteristics**

| | Density of mixed cell clusters | | |
|---|---|---|---|
| | **Spearman r** | **95% confidence interval** | **P value** |
| Age (yrs.) | 0,03 | [-0.34 to 0.39] | 0,88 |
| Body-mass index ($kg/m^2$) | -0,02 | [-0.38 to 0.35] | 0,93 |
| Pack years (no.) | 0,23 | [-0.16 to 0.55] | 0,24 |
| **LFT** | | | |
| $FEV_1$ (% pred.) | -0,39 | [-0.66 to -0.03] | **0,03** |
| $FEV_1$/FVC ratio (%) | -0,36 | [-0.65 to 0.009] | **0,04** |
| FVC (% pred.) | -0,35 | [-0.63 to 0.02] | 0,057 |
| RV (% pred.) | - 0,29 | [-0.08 to 0.59] | 0,11 |
| TLCO (% pred.) | -0,48 | [-0.73 to -0.12] | **0,008** |
| **Six-minute walk test distance (m)** | 0,11 | [-0.33 to 0.51] | 0,62 |
| **Arterial blood gases** | | | |
| $PaO_2$ (mm Hg) | 0,21 | [-0.18 to 0.54] | 0,28 |
| $PaCO_2$ (mm Hg) | 0,33 | [-0.06 to 0.62] | 0,08 |
| **CT parameters** | | | |
| Bronchi: | | | |
| WA4% | 0,14 | [-0.29 to 0.51] | 0,52 |
| WT4 (mm) | 0,13 | [-0.29 to 0.51] | 0,54 |
| WA5% | 0,24 | [-0.19 to 0.59] | 0,25 |
| WT5 (mm) | 0,32 | [-0.09 to 0.64] | 0,12 |
| Emphysema: | | | |
| LAA (%) | 0,30 | [-0.09 to 0.61] | 0,12 |
| Air trapping: | | | |
| MLA E (HU) | -0,22 | [-0.59 to 0.22] | 0,30 |
| MLA I (HU) | -0,38 | [-0.66 to 0.006] | **0,047** |
| MLA I-E (HU) | 0,11 | [-0.32 to 0.51 | 0,60 |
| Pulmonary Vessels | | | |
| $\%CSA_{<5}$ | -0,25 | [-0.58 to 0.13] | 0,18 |
| $\%CSA_{5-10}$ | -0,28 | [-0.59 to 0.11] | 0,15 |
| $CSN_{<5}$ | -0,20 | [-0.54 to 0.19] | 0,29 |
| $CSN_{5-10}$ | -0,39 | [-0.66 to -0.01] | **0,04** |

$FEV_1$, forced expiratory volume in 1 second; FVC, forced vital capacity; LFT, lung function test; RV, residual volume; TLCO, Transfer Lung capacity of Carbon monoxide, $PaO_2$, partial arterial oxygen pressure, $PaCO_2$, partial arterial carbon dioxide pressure; WA, mean wall area; LA, mean lumen area, WA%, mean wall area percentage; WT, wall thickness; LAA, low-attenuation area; MLA E or I, mean lung attenuation value during expiration or inspiration. MLA I-E, difference between inspiratory and expiratory mean lung attenuation value. $\%CSA_{<5}$, percentage of total lung area taken up by the cross-sectional area of pulmonary vessels less than 5 $mm^2$; $\%CSA_{5–10}$, percentage of total lung area taken up by the cross-sectional area of pulmonary vessels between 5 and 10 $mm^2$; $CSN_{<5}$, number of vessels less than 5 $mm^2$ normalized by total lung area; $CSN_{5-10}$, number of vessels between 5 and 10 $mm^2$ normalized by total lung area; NR: not relevant. The correlation coefficient (r), 95% confidence interval and significance level (P value), were obtained by using nonparametric Spearman analysis.

**Table S6. Multivariate analysis of $FEV_1$/FVC**

| $FEV_1$/FVC | | | | | |
|---|---|---|---|---|---|
| **Model** | **Explicative variables** | **Coefficient** | **Standard error** | **T-value** | **P value** |
| $R^2$= 0.39,<br>Adjusted $R^2$= 0.35<br>P = 0.0009<br>F = 9.04<br>Residual standard error = 0.12 | Interacting cells density<br><br>Density of mixed cells clusters | -0.0025<br><br>0.00230 | 0.0006<br><br>0.0011 | -4.0<br><br>2.2 | 0.0004<br><br>0.04 |

$FEV_1$, forced expiratory volume in 1 second; FVC, forced vital capacity

**Table S7. Patient characteristics (for tissular CD8$^+$ T cells purification)**

| | COPD | Control |
|---|---|---|
| n | 20 | 26 |
| Age (yr) | 64.0 ± 9.5 | 65.7 ± 12.5 |
| Sex (Men/Woman) | 12/8 | 11/15 |
| Body-mass index (kg/m$^2$) | 23.8 ± 4.0 | 25.9 ± 4.2 |
| Current smoker (Y/N) | 6/14 | 0/26 |
| Former smoker (Y/N) | 14/6 | 19/7 |
| Pack years (no.) | 48.2 ± 26.9 | 24.0 ± 21.7 |
| **PFT** | | |
| $FEV_1$ (% pred.) | 60.7 ± 27.0 | 98.1 ± 20.2 |
| $FEV_1$/FVC ratio (%) | 55.1 ± 13.7 | 79.3 ± 6.4 |

Plus–minus values are means ± SD. PFT, pulmonary function test; $FEV_1$, forced expiratory volume in 1 second; FVC, forced vital capacity.

**Table S8: Patient characteristics (for circulating $CD8^+$/$CD4^+$ T cells and fibrocyte precursors purification)**

| | COPD |
|---|---|
| n | 44 |
| Age (years) | 67.4 ± 7.7 |
| Sex (Men/Woman) | 24/20 |
| Body-mass index ($kg/m^2$) | 29.2 ± 7.0 |
| Current /Former smokers | 16/28 |
| Pack years (no.) | 48.1 ± 19.3 |
| **PFT** | |
| $FEV_1$ (% pred.) | 59.7 ± 18.0 |
| $FEV_1$/FVC ratio (%) | 56.9 ± 11.5 |
| FVC (% pred.) | 83.3 ± 15.4 |
| **Six-minute walk test distance (m)** | 450 ± 133 |
| **Arterial blood gases** | |
| $PaO_2$ (mm Hg) | 73.7 ± 11.2 |
| $PaCO_2$ (mm Hg) | 38.8 ± 5.3 |

Plus–minus values are means ± SD. PFT, pulmonary function test; $FEV_1$, forced expiratory volume in 1 second; FVC, forced vital capacity; $PaO_2$, partial arterial oxygen pressure, $PaCO_2$, partial arterial carbon dioxide pressure.

**Table S9: Patient characteristics (for basal bronchial epithelial cell purification)**

| | Patients |
|---|---|
| n | 2 |
| Age (yr) | 64.1 ± 9.6 |
| Sex (Men/Woman) | 0/2 |
| Body-mass index (kg/m$^2$) | 19.0 ± 3.4 |
| Current smoker (Y/N) | 1/1 |
| Former smoker (Y/N) | 1/1 |
| Pack years (no.) | 22.5 ± 24.7 |
| **PFT** | |
| $FEV_1$ (% pred.) | 90.4 ± 20.3 |
| $FEV_1$/FVC ratio (%) | 70.0 ± 7.7 |

Plus–minus values are means ± SD. PFT, pulmonary function test; $FEV_1$, forced expiratory volume in 1 second; FVC, forced vital capacity.

**Table S10. Definition of the notations and parameters of the mathematical model**

| | Symbol | Meaning |
|---|---|---|
| **General** | $L$ | Lamina propria (=peribronchial area) |
| | $x_0$ | Side length of the units of the lattice $L$ |
| | $M(s)$ | Neighbourhood of the site $(s)$ |
| | $V(s)$ | Number of F and C cells belonging to $M(s)$ |
| | $V(F)(s)$ | Number of F cells belonging to $M(s)$ |
| | $V(C)(s)$ | Number of C cells belonging to $M(s)$ |
| | $N_k(F)$ | Number of F cells at the beginning of period $k$ |
| | $N_k(C)$ | Number of C cells at the beginning of period $k$ |
| **Initial situation** | $n_0(C)$ | Initial density of C cells |
| | $n_0(F)$ | Initial density of F cells |
| | $N_0(C)$ | Initial number of C cells |
| | $N_0(F)$ | Initial number of F cells |
| **Cell death** | $p_{dF}$ | Probability for a F cell to die |
| | $p_{dC}$ | Basal probability for a C cell to die |
| | $p_{dC+}$ | Increased probability for a C cell to die |
| | σ | Threshold number of neighbouring C cells, above which the probability of dying is increased from $p_{dC}$ to $p_{dC+}$ |
| **Cell proliferation** | $p_F$ | Probability for a F cell to divide |
| | $p_C$ | Basal probability for a C cell to divide |
| | $p_{C/F}$ | Increased probability for a C cell to divide |
| | λ | Threshold number of neighbouring C cells of an empty s' site belonging to M(s), above which the considered C cell does not divide. |
| **Cell displacement** | $P_F\,(s,s')$ | Probability for a F cell to go from s to s' |
| | $P_C\,(s,s')$ | Probability for a C cell to go from s to s' |
| | $f_F$ | Function partially defining $P_F\,(s,s')$ when $s' \in$ M(s), $s'$ is empty and $s' \neq s$, depending on $V(C)(s')$ |
| | $f_C$ | Function partially defining $P_C\,(s,s')$ when $s' \in$ M(s), $s'$ is empty and $s' \neq s$, depending on $V(s')$ |
| | $\varepsilon_F$ | Value taken by $f_F$ to reflect a low attraction |
| | $\varepsilon_C$ | Value taken by $f_C$ to reflect a low attraction |
| **Cell infiltration** | $p_{istaF}$ | Probability for a F cell to get infiltrated at the beginning of a 3 minutes-period |
| | $p_{istaC}$ | Probability for a C cell to get infiltrated at the beginning of a 3 minutes-period |

| | | |
|---|---|---|
| | $p_{iexaF}$ | Probability for a F cell to get infiltrated during an exacerbation |
| | $p_{iexaC}$ | Probability for a F cell to get infiltrated during an exacerbation |
| | $N_{iexaF}$ | Number of F cells that are infiltrated during an exacerbation |
| | $N_{iexaC}$ | Number of C cells that are infiltrated during an exacerbation |

**Table S11**. Numerical values of parameters depending in control and COPD situations

| Symbol | Numerical values | |
|---|---|---|
| | **Control** | **COPD** |
| $x_0$ | 7 µm | |
| $N_0(C)$ | 118 cells | |
| $N_0(F)$ | 19 cells | |
| $p_{dF}$ | $4.8.10^{-6}$ | $2.4.10^{-6}$ |
| $p_{dC}$ | $1.10^{-4}$ | $5.10^{-5}$ |
| $p_{dC+}$ | $4.10^{-4}$ | $2.10^{-4}$ |
| σ | 3 | |
| $p_F$ | 0 | |
| $p_C$ | $5.10^{-5}$ | |
| $p_{C/F}$ | $2.10^{-4}$ | |
| λ | 3 | |
| $\varepsilon_F$ | $10^{-3}$ | |
| $\varepsilon_C$ | $10^{-3}$ | |
| $p_{istaF}$ | $9.12.10^{-5}$ | |
| $p_{istaC}$ | $1.40.10^{-2}$ | |
| $p_{iexaF}$ | 0 | $2.20.10^{-3}$ |
| $p_{iexaC}$ | 0 | |
| $N_{iexaF}$ | 0 | 1 |
| $N_{iexaC}$ | 0 | |

**Movie S1 (separate file).** Two day after adding non-activated CD8+ T cells (bright round cells) on fibrocytes (adherent elongated cells), phase-contrast images of co-culture taken were recorded every 2 minutes. A tracked lymphocyte is indicated by a blue dot and its trajectory is shown by a blue line dot (Manual Tracking plugin, Fiji software).

**Movie S2 (separate file). Cell dynamics within the peribronchial area, 2 years after the initial time, with control dynamics**. Images of the simulations were recorded every 3 min during 24 hours. $CD8^+$ T cells and fibrocytes are represented respectively by pink and green squares. control (resp. COPD) situation.

**Movie S3 (separate file). Cell dynamics within the peribronchial area, 2 years after the initial time, with COPD dynamics**. Images of the simulations were recorded every 3 min during 24 hours. $CD8^+$ T cells and fibrocytes are represented respectively by pink and green squares. control (resp. COPD) situation.

**Movie S4 (separate file). Cell dynamics within the peribronchial area, 7 years after the initial time, with control dynamics**. Images of the simulations were recorded every 3 min during 24 hours. $CD8^+$ T cells and fibrocytes are represented respectively by pink and green squares. control (resp. COPD) situation.

**Movie S5 (separate file). Cell dynamics within the peribronchial area, 7 years after the initial time, with COPD dynamics**. Images of the simulations were recorded every 3 min during 24 hours. $CD8^+$ T cells and fibrocytes are represented respectively by pink and green squares. control (resp. COPD) situation.
